# *b3galt6* knock-out zebrafish recapitulate β3GalT6-deficiency disorders in human and reveal a trisaccharide proteoglycan linkage region

**DOI:** 10.1101/2020.06.22.165316

**Authors:** Sarah Delbaere, Adelbert De Clercq, Shuji Mizumoto, Fredrik Noborn, Jan Willem Bek, Lien Alluyn, Charlotte Gistelinck, Delfien Syx, Phil L. Salmon, Paul J. Coucke, Göran Larson, Shuhei Yamada, Andy Willaert, Fransiska Malfait

**Affiliations:** Center for Medical Genetics Ghent, Department of Biomolecular Medicine, Ghent University and Ghent University Hospital, Ghent, Belgium; Department of Pathobiochemistry, Faculty of Pharmacy, Meijo University, 150 Yagotoyama, Tempaku-ku, Nagoya, Aichi 468-8503, Japan; Department of Laboratory Medicine, Sahlgrenska Academy at the University of Gothenburg, Gothenburg, Sweden; Laboratory of Clinical Chemistry, Sahlgrenska University Hospital, Gothenburg, Sweden; Department of Orthopaedics and Sports Medicine, University of Washington, Seattle, WA98195; Bruker microCT, Kontich, Belgium

**Author notes:** AW and FM contributed equally to this work. Corresponding author: Fransiska Malfait, Center for Medical Genetics, Ghent University Hospital, 0K5, Corneel Heymanslaan 10, B-9000 Ghent, Belgium. Author Contributions SD, AW and FM designed the study; SD, ADC, SM, FN, JWB, LA and CG performed research; SM, PLS, GL, SY contributed new reagents or analytic tools; SD, ADC, SM, FN, JWB, LA, CG, DS, PLS, PJC, GL, SY, AW and FM analyzed data; SD, AW and FM wrote the paper; all authors reviewed the paper.

**Keywords:** *b3galt6*, zebrafish, trisaccharide linkage region, proteoglycans, linkeropathies

## Abstract

Proteoglycans are structurally and functionally diverse biomacromolecules found abundantly on cell membranes and in the extracellular matrix. They consist of a core protein linked to glycosaminoglycan chains via a tetrasaccharide linkage region. Here, we show that CRISPR/Cas9-mediated *b3galt6* knock-out zebrafish, lacking galactosyltransferase II, which adds the third sugar in the linkage region, largely recapitulate the phenotypic abnormalities seen in human β3GalT6-deficiency disorders. These comprise craniofacial dysmorphism, generalized skeletal dysplasia, skin involvement and indications for muscle hypotonia. In-depth TEM analysis revealed disturbed collagen fibril organization as the most consistent ultrastructural characteristic throughout different affected tissues. Strikingly, despite a strong reduction in glycosaminoglycan content, as demonstrated by anion-exchange HPLC, subsequent LC-MS/MS analysis revealed a small amount of proteoglycans containing a unique linkage region consisting of only three sugars. This implies that formation of glycosaminoglycans with an immature linkage region is possible in a pathogenic context. Our study therefore unveils a novel rescue mechanism for proteoglycan production in the absence of galactosyltransferase II, hereby opening new avenues for therapeutic intervention.

## Introduction

Proteoglycans (PGs) comprise a group of complex biomacromolecules, consisting of a core protein and one or multiple glycosaminoglycan (GAG) side chains, that are ubiquitously expressed in the extracellular matrix (ECM) and on cell surfaces and basement membranes in vertebrate and invertebrate species [1-4]. Besides their structural role, PGs act as mediators between the ECM and intracellular signaling pathways and contribute to development and tissue homeostasis, influencing a variety of cellular processes including cell fate determination, cell proliferation, migration, adhesion, differentiation and survival [5].

The PG superfamily is, depending on the composition of the GAG chains, subdivided into heparan sulfate (HS) and chondroitin/dermatan sulfate (CS/DS) PGs. These GAG chains are linear polysaccharides, composed of a repeated disaccharide unit consisting of an uronic acid and amino sugar. HSPGs, such as perlecan, syndecan and glypican, consist of repeating glucuronic acid (GlcA)-*N*-acetyl-glucosamine (GlcNAc) disaccharides. CSPGs consist of repeating GlcA-*N*-acetyl-galactosamine (GalNAc) disaccharides. DS is a stereoisomer of CS, in which GlcA is epimerized to iduronic acid (IdoA). In mammalian tissues, CS and DS chains are often present as hybrid CS-DS chains on PGs (e.g. decorin, biglycan) [6]. Tightly controlled modifications, such as epimerization and sulfation reactions, further increase the structural and functional diversity of the HS and CS/DS PGs [7, 8]. GAG biosynthesis of both HSPGs and CS/DSPGs is initiated in the endoplasmic reticulum and Golgi apparatus by the formation of a common tetrasaccharide linkage region (glucuronic acid-galactose-galactose-xylose-*O*--) (GlcA-Gal-Gal-Xyl-*O*-) via *O*-glycosylation of a serine residue of the core protein (Fig. 1A) [1-3]. The first step in the biosynthesis of this tetrasaccharide linkage region is catalyzed by xylosyltransferases (XylTs) encoded by the paralogues *XYLT1* [MIM 608124] and *XYLT2* [MIM 608125] [9]. Two Gal residues are subsequently added by respectively galactosyltransferase I (GalT-I or β4GalT7 encoded by *B4GALT7* [MIM 604327]) and galactosyltransferase II (GalT-II or β3GalT6 encoded by *B3GALT6* [MIM 615291]) [10, 11]. Further addition of a GlcA by glucuronosyltransferase I (GlcAT-I encoded by *B3GAT3)* [MIM 606374] completes the formation of the linkage region [12].

**Fig. 1.**
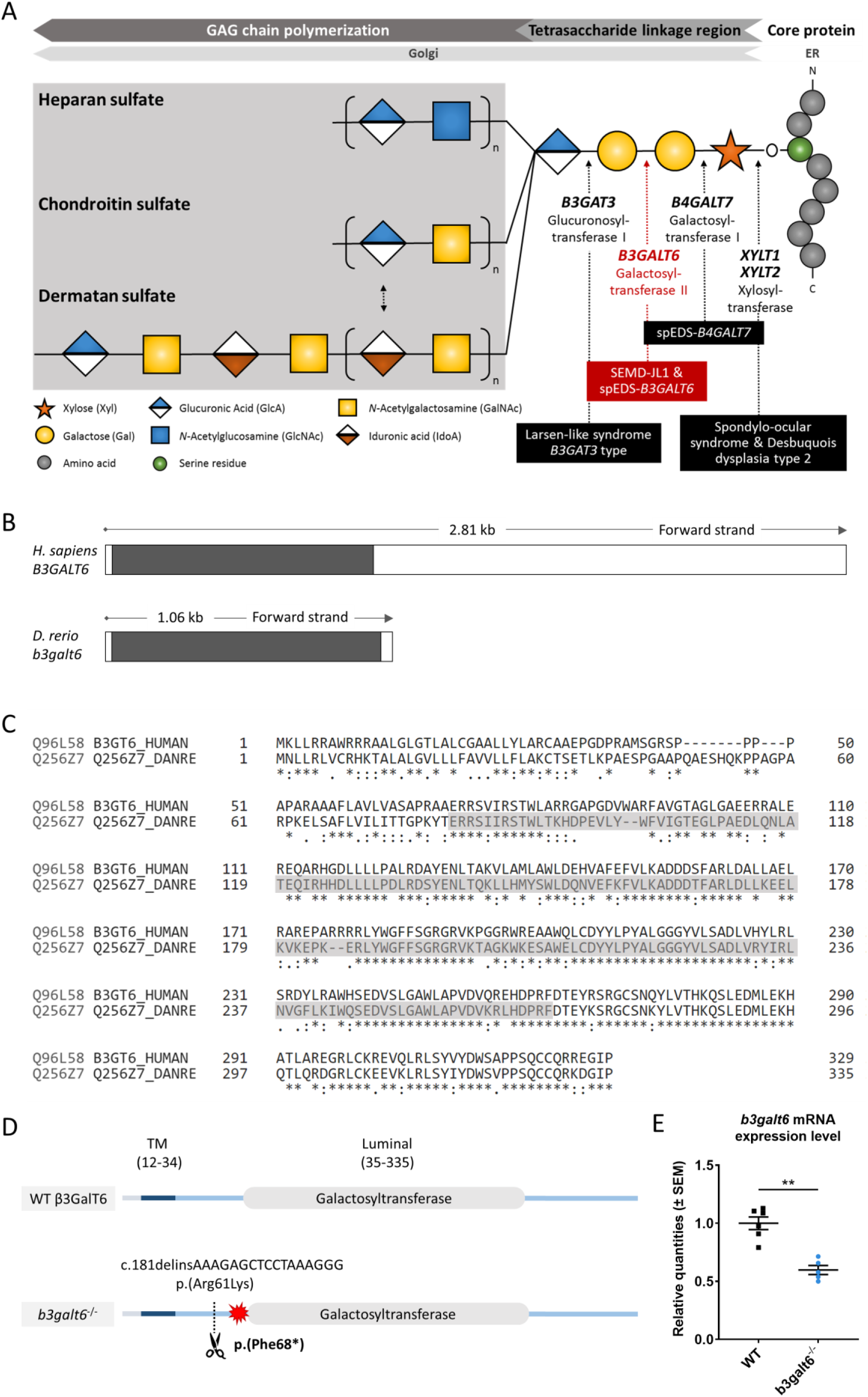
Schematic representation of the GAG biosynthesis and the evolutionary conservation of the zebrafish and human *b3galt6*/*B3GALT6* gene and B3galt6/β3GalT6 protein. (**A**) Simplified schematic representation of the PG linkage and glycosaminoglycans biosynthesis. PG core proteins are synthesized in the endoplasmic reticulum and undergo further modification in the Golgi apparatus. First, a tetrasaccharide linkage region is synthesized that originates from the addition of a Xyl onto a serine residue of the core protein. Subsequent addition of two Gals and one GlcA residues completes this process. Depending on the addition of GlcNAc or GalNAc to the terminal GlcA residue of the linkage region, HS or CS/DS PGs will be formed, respectively. The elongation of the GAG chain continues by repeated addition of uronic acid and amino sugar residues. The GAG chain is then further modified by epimerization and sulfation (not shown). The main genes encoding for the enzymes involved in the linkage region biosynthesis are indicated in bold and biallelic mutations in these genes result in different linkeropathies. Disorders resulting from deficiency of the enzymes are indicated in black boxes. The deficient enzyme (β3GalT6) and the related disorder studied here are depicted in red (e.g. spEDS-*B3GALT6* and SEMD-JL1). (**B**) Schematic representation of the genomic structure of the *H. sapiens B3GALT6* (NM_080605.4) and *D. rerio b3galt6* (NM_001045225.1) genes. The white and black boxes represent the untranslated regions and the single *B3GALT6*/*b3galt6* exon, respectively. The zebrafish *b3galt6* gene resides on chromosome 11, spans a region of 1.06 kb and has not been duplicated during teleost evolution. (**C**) Amino acid (AA) sequence alignment (Clustal Omega) between *H. sapiens* β3GalT6 (Uniprot: Q96L58) and *D. rerio* B3galt6 (Uniprot: Q256Z7). The luminal galactosyltransferase domain is highlighted in grey. Conservation of AA sequences is shown below the alignment: “*” residues identical in all sequences in the alignment; “:” conserved substitutions; “.” semi-conserved substitutions; “space” no conservation. (**D**) Schematic representation of the cmg20 allele in *b3galt6* mutants. *b3galt6* consists of three parts, a cytosolic domain (AA 1-11), a transmembrane domain (TM, AA 12-34) and a luminal domain (AA 35-335). The location of the CRISPR/Cas9-generated indel variant is indicated by a scissor symbol and the resulting premature termination codon (PTC) is depicted by a red star. (**E**) RT-qPCR analysis of relative *b3galt6* mRNA expression levels in *b3galt6*^−/−^ adult zebrafish (0.597 ± 0.03919) (n=5) compared to WT siblings (1 ± 0.05395) (n=6). Data are expressed as mean ± SEM. A Mann-Whitney U test was used to determine significance, **P<0.01.

The cellular machinery required for GAG biosynthesis and modification is conserved over a broad range of eukaryotic organisms [13]. The particular importance of a correct initiation of GAG synthesis is underscored by the identification of a series of severe, overlapping and multisystemic genetic human disorders that result from deficient activity of any of these “linkage-enzymes”, collectively coined as ‘linkeropathies’ [14-16]. The most frequently reported linkeropathy results from galactosyltransferase II deficiency, caused by biallelic pathogenic variants in *B3GALT6*. This leads to two clinically overlapping conditions, spondylodysplastic Ehlers-Danlos syndrome (EDS) (spEDS-*B3GALT6*) and spondyloepimetaphyseal dysplasia with joint laxity type I (SEMD-JL1) [16, 17], both of which are characterized by variable degrees of spondyloepimetaphyseal bone dysplasia, (with postnatal growth restriction, bowing of long bones, hypo- or dysplasia of iliac and long bones, and vertebral body changes), kyphoscoliosis, bone fragility with spontaneous fractures, joint hypermobility in conjunction with joint contractures, hypotonia with delayed motor development, skin fragility, and craniofacial dysmorphisms, including midfacial hypoplasia and dysplastic teeth [16-18]. The severe and pleiotropic phenotype suggests a critical role for *B3GALT6* in the development and homeostasis of various connective tissues. Pathogenic variants in *B3GALT6* either result in mislocalization and/or reduced amounts of active galactosyltransferase II [19]. Previous studies on several biallelic *B3GALT6* variants showed that they resulted in a barely detectable *in vitro* galactosyltransferase II enzymatic activity, but intriguingly HS and CS/DS GAG synthesis was partially preserved *in cellulo* [20]. While some variants may have a greater deleterious effect on the *in vitro* than on the *in cellulo* enzymatic activity, it is currently unknown whether there are rescue mechanisms at play to (partially) compensate for the galactosyltransferase II activity.

In recent years, zebrafish have emerged as an excellent tool to model disease and to increase our understanding of developmental processes and disease mechanisms. Zebrafish share 70% of their genes with humans [21] and the advantages of their use are reflected by their large breed size, short lifecycle and low husbandry costs [22]. Zebrafish has proven to be a relevant genetic model for the study of PG biosynthesis since many enzymes involved in this process are strongly conserved between human and zebrafish [23], and a number of zebrafish models for defective PG synthesis (including *fam20b*^*b1125*^, *xylt1*^*b1189*^, *or b3gat3*^*hi307*^), provided novel knowledge on PG function in bone and cartilage formation [23-26].

In this study, we generated *b3galt6* knock-out (KO) zebrafish as a model for spEDS-*B3GALT6* and SEMD-JL1. These models were used to investigate the structure and amount of different GAG disaccharides and the PG linkage region composition in these disorders. Furthermore, in-depth phenotypic characterization of *b3galt6*^*−/−*^ zebrafish was done to reveal to which extent key features of the human disorders are recapitulated and to further elucidate the function of galactosyltransferase II in vertebrate musculoskeletal development.

## Results

### CRISPR/Cas9-mediated *b3galt6* knock-out in zebrafish leads to reduced HS and CS/DS glycosaminoglycan concentrations in bone, muscle and skin

Similar to its human orthologue, zebrafish *b3galt6* (NM_001045225.1) is a single-exon gene (Fig. 1B). Human and zebrafish galactosyltransferase II (β3GalT6/B3galt6) protein sequence display 82.5 % amino acid (AA) similarity, with the highest AA conservation in the luminal domain (Fig. 1C).

In order to investigate the consequences of zebrafish B3galt6-deficiency on GAG biosynthesis in different tissues, we generated two *b3galt6* KO zebrafish models using CRISPR/Cas9 gene editing. The first KO model carries the c.181delinsAAAGAGCTCCTAAAGGG indel mutation (cmg20 allele), leading to a premature termination codon (PTC) (p.(Phe68*)), located upstream to the catalytic domain (Fig. 1D). RT-qPCR showed a 40.3% reduction of *b3galt6 mRNA* expression in homozygous adult *b3galt6* KO zebrafish compared to wild-type (WT) siblings (Fig. 1E). For the remainder of this manuscript, this model will be referred to as *b3galt6*^−/−^. The second KO model carries a four base pair deletion (c.398_401del) (cmg22 allele), predicted to generate a PTC (p.(Asn139*)) in the catalytic domain (NM_001045225.1). This mutant was solely used for validation purposes (Fig. S1). Both *b3galt6* KO zebrafish were produced at an expected Mendelian ratio and survived into adulthood. Unfortunately, commercial antibodies, which were only available against human β3GalT6, did not work in our hands and precluded the investigation of zebrafish B3galt6 protein levels and localization.

To examine the activity of B3galt6 in different *b3galt6*^−/−^ tissues, concentrations of CS, DS and HS disaccharides extracted from bone, muscle and skin from WT and *b3galt6*^−/−^ zebrafish were quantified using a combination of enzymatic digestion and anion-exchange HPLC (Figs. S2-S5 and Tables S1-S4). Overall, a generalized decrease in HS, CS and DS concentration was detected in all examined tissues of *b3galt6*^−/−^ zebrafish. In the GAG fractions prepared from bone, the disaccharide concentration of the CS/DS and CS moieties was significantly decreased by respectively 55% and 64%, compared to WT zebrafish (Table 1 and Tables S1 and S2). DS moieties were undetectable in bone of *b3galt6*^−/−^ zebrafish, whereas their concentration was 50.5 pmol/mg protein in bone of WT zebrafish (Table 1 and Table S3). A non-significant decrease in the disaccharide concentration of the HS moiety was observed in *b3galt6*^−/−^ mutant bone (Table 1 and Table S4), compared to WT. In GAG fractions prepared from *b3galt6*^−/−^ muscle, CS/DS, CS and DS disaccharide moieties were generally lower than in WT muscle (Table 1 and Tables S1-S3). Due to the low concentrations close to the detection limit, a significant reduction could only be detected for the DS moiety (72% reduction) (Table 1 and Table S3). In addition, a significant decrease in HS disaccharide moieties by 56% was noted in *b3galt6*^−/−^ muscle compared to WT muscle (Table 1 and Table S4). In the GAG fractions prepared from *b3galt6*^*−/−*^ skin, the concentration of all disaccharide moieties was significantly reduced compared to WT (Table 1 and Tables S1-S4). The disaccharide concentration of the CS/DS, CS, DS and HS moieties were decreased by respectively 77%, 73%, 55% and 62% compared to WT (Table 1 and Tables S1-S4). No altered sulfation patterns were observed in disaccharide moieties when comparing unique GAG chains between WT and *b3galt6*^−/−^ zebrafish samples (Tables S1-S4).

**Table 1.** Concentrations of GAG disaccharides in wild-type and *b3galt6*^−/−^ zebrafish tissues

|  | *pmol/mg protein (Average ± SE)* |  |  |  |
| --- | --- | --- | --- | --- |
|  | CS/DS | CS | DS | HS |
| **Bone** |  |  |  |  |
| Wild-type | 774.2 ± 19.9 | 728.4 ± 148.1 | 50.5 ± 8.1 | 178.6 ± 19.8 |
| *b3galt6*<sup>-/-</sup> | 346.1 ± 20.0* | 279.9 ± 4.8** | N.D. | 137.1 ± 39.7<sup>ns</sup> |
| **Muscle** |  |  |  |  |
| Wild-type | 9.3 ± 0.4 | 2.1 ± 0.4 | 14.5 ± 1.1 | 30.8 ± 1.2 |
| *b3galt6*<sup>-/-</sup> | 5.7 ± 3.1<sup>ns</sup> | N.D. | 4.1 ± 1.0* | 13.5 ± 4.0** |
| **Skin** |  |  |  |  |
| Wild-type | 236.2 ± 26.5 | 175.1 ± 25.3 | 75.9 ± 7.9 | 55.1 ± 4.5 |
| *b3galt6*<sup>-/-</sup> | 55.4 ± 15.2* | 47.9 ± 13.5** | 33.8 ± 6.3** | 21.0 ± 2.8** |

Student's t-test (vs wild-type) (n=3), N.D.: not detected (<1 pmol/mg protein), \*P<0.01, \*\*P<0.05, ns: not significant, SE: standard error.

Despite the strong reduction in disaccharide concentrations observed in *b3galt6*^−/−^ zebrafish, low amounts of GAGs were still produced in the mutants. Similar results were obtained for the second *b3galt6* KO zebrafish (cmg22 allele) (Table S5). RT-qPCR revealed no significant upregulation of other galactosyltransferase(s) member(s) from the *b3galt* family (*b3galt1a, b3galt2, b3galt4, b3galnt2*) or of other linkage-enzymes (*xylt1, xylt2, b4galt7* and *b3gat3*) (Fig. S6), suggesting that altered expression of these genes did not compensate for the reduced *b3galt6* mRNA levels.

### B3galt6-deficiency leads to the biosynthesis of a non-canonical (GlcA-Gal-Xyl-*O*-) trisaccharide linkage region in zebrafish

In order to explain why *b3galt6*^−/−^ zebrafish can still produce GAG chains on PG core proteins, we investigated the composition of the linkage region. To this purpose, whole zebrafish extracts were enriched for all proteoglycans, their GAG chains depolymerized with bacterial lyases, the core proteins digested with trypsin and the remaining glycopeptides subjected to nano-scale liquid chromatography-tandem mass spectrometry (nLC-MS/MS) using a recently developed glycoproteomic approach [27, 28]. Potential structural differences in PG linkage regions in WT versus *b3galt6*^−/−^ zebrafish are thus elucidated through the combined identities of peptide backbones and GAG linkage region structures. The glycoproteomic analysis identified two CS-glycopeptides derived from biglycan (UniProt: A8BBH0) in both the WT and *b3galt6*^−/−^ zebrafish. These CS-glycopeptides displayed different linkage region structures in WT compared with *b3galt6*^−/−^ zebrafish (Fig. 2). While the peptide in the WT zebrafish was modified with a hexasaccharide structure (Fig. 2A), the peptide in the *b3galt6*^−/−^ zebrafish was modified with a pentasaccharide structure (Fig. 2B), supporting the hypothesis that *b3galt6*^−/−^ zebrafish produces a non-canonical trisaccharide linkage region. The precursor ion (m/z 1281.7922; 3+) in the WT zebrafish equated to a mass of a peptide (DQEEGSAVEPYKPEHPTCPFGCR) modified with a hexasaccharide structure, with the peptide containing two carbamidomethyl (CAM) modifications. Furthermore, the hexasaccharide structure was modified with one sulfate group (SO_3_^−^) on the subterminal GalNAc residue [GlcAGalNAc-H_2_O+SO_3_+H]^+^, and one phosphate group (HPO_3_^−^) on the Xyl residue (Fig. S7). In contrast, the precursor ion (*m/z* 1227.7765; 3+) in the *b3galt6*^−/−^ zebrafish equated to a mass of the same peptide but modified with a pentasaccharide structure, with two phosphate/sulfate groups, with the peptide containing two CAM modifications. The difference in mass between the WT and *b3galt6*^−/−^ precursor ions corresponds to 162.08 Da, equating to the mass of a hexose (galactose) residue (162 Da). Additional HS- and CS-glycopeptides with the expected tetrasaccharide linkage region were found in the WT. In the *b3galt6*^−/−^, no HS-modified structures were identified but additional CS-glycopeptides with the non-canonical trisaccharide linkage region were found. Table S6 shows a complete list of all glycopeptides identified in both WT and *b3galt6*^−/−^ zebrafish. PGs harboring a tetrasaccharide linkage region were never detected in the *b3galt6*^−/−^ adult zebrafish samples and no trisaccharide linkage region was detected in WT zebrafish samples.

**Fig. 2.**
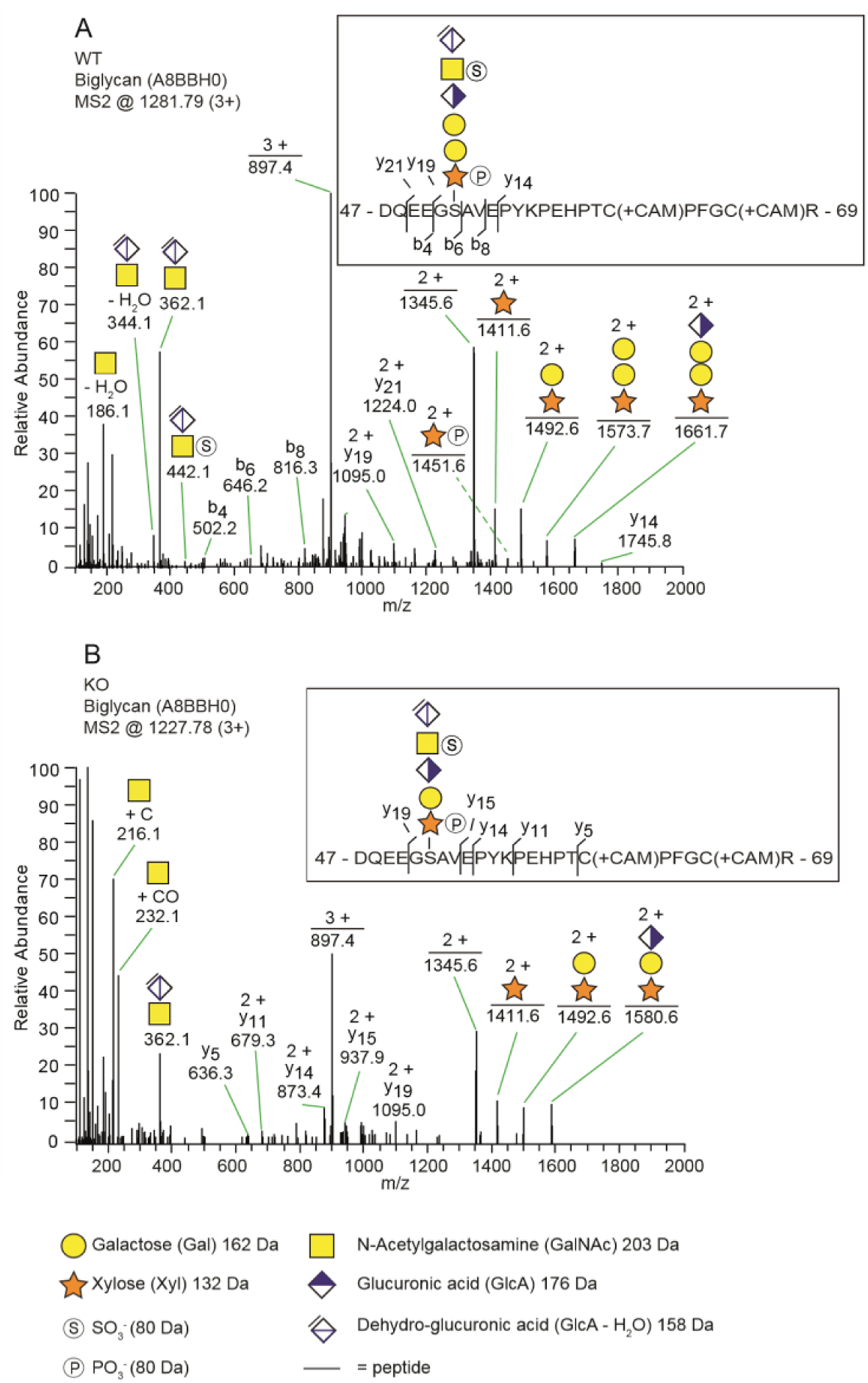
The identification of a trisaccharide linkage region in the *b3galt6*^−/−^ zebrafish. (**A** and **B**) MS2 fragment mass spectra of CS-glycopeptides derived from biglycan (UniProt: A8BBH0) showed different linkage regions in WT compared to the *b3galt6*^−/−^ zebrafish. (**A**) The peptide (DQEEGSAVEPYKPEHPTCPFGCR) in WT zebrafish was modified with a residual hexasaccharide structure, corresponding to the canonical tetrasaccharide linkage region (*m/z* 1281.79; 3+). The hexasaccharide structure was modified with one sulfate group (SO_3_^−^) at the subterminal GalNAc residue and one phosphate group (HPO_3_^−^) at the xylose residue (Fig. S7) (**B**) The peptide (DQEEGSAVEPYKPEHPTCPFGCR) in *b3galt6*^−/−^ zebrafish was modified with a residual pentasaccharide structure, corresponding to a non-canonical trisaccharide linkage region (*m/z* 1227.78; 3+). The pentasaccharide structure was modified with two phosphate/sulfate groups. Their exact positioning and identity could not be determined due to low intensities of corresponding ion peaks, and the positions are therefore assigned in analogy of the WT structure.

These observations suggest that B3GAT3/GlcAT-I can transfer a GlcA residue not only to Gal-Gal-Xyl, but also to a Gal-Xyl substrate. This was confirmed by an *in vitro* GlcAT-I assay, which showed that GlcAT-I is indeed capable of adding a GlcA to a Gal-Xyl-*p*-nitrophynyl, which was chemically synthesized, thus to a substrate ending with a single galactose (Fig. S8).

### *b3galt6*^−/−^ zebrafish show early-onset, progressive morphological abnormalities and delay of cartilage and bone formation, reminiscent of human spEDS-*B3GALT6*

One of the clinical hallmarks of human spEDS-*B3GALT6* is the pronounced skeletal involvement with the presence of spondyloepimetaphyseal dysplasia with postnatal growth restriction, bowing of long bones, hypoplasia of iliac bones, femoral head dysplasia, vertebral body changes, kyphoscoliosis and bone fragility with spontaneous fractures (Table 2). To evaluate the development of cartilage and bone associated with B3galt6-deficiency in zebrafish, phenotypic characterization of *b3galt6*^*−/−*^ zebrafish was performed in juvenile (20 dpf, days post fertilization) and adult stages (four months).

**Table 2.** Clinical features and ultrastructural and biochemical abnormalities observed in spEDS-*B3GALT6* and SEMD-JL1 individuals compared to the *b3galt6*^−/−^ zebrafish model. n.a.: not available. GAG: glycosaminoglycan. The columns ‘spEDS-*B3GALT6*’ and ‘SEMD-JL1’ summarize the most important clinical features of all hitherto reported patients for whom this data was available [20, 47].

|  | Human |  | Zebrafish |
| --- | --- | --- | --- |
|  | spEDS-*B3GALT6* | SEMD-JL1 | *b3galt6*<sup>-/-</sup> mutant |
| General morphology |  |  |  |
| Short stature | 22/30 | 11/14 | Reduced body length |
| Craniofacial dysmorphisms (e.g. midfacial hypoplasia, micrognathia) | 25/30 | 12/14 | Deformed, small and round head with shorter parietal/occipital region and underdeveloped olfactory region |
| Dental abnormalities | 14/30 | n.a. | Smaller teeth |
| Musculoskeletal involvement |  |  |  |
| Bowing of limbs, metaphyseal widening or vertebral body changes | 20/30 | 14/14 | Deformities of the fin skeleton |
| Bone fragility with spontaneous fractures | 19/30 | 0/3 | Extra intramembranous bone, lower degree of collagen type I fibril organization and higher TMD after correction for smaller size |
| Joint contractures (especially hands) | 23/30 | 5/14 | Pectoral fins point upwards |
| Joint hypermobility | 26/30 | 6/14 | n.a. |
| Kyphoscoliosis | 23/30 | 13/14 | Kyphosis and scoliosis |
| Muscle hypotonia | 18/30 | 1/14 | Reduced critical swimming speed and endurance |
| Skin features |  |  |  |
| Skin hyperextensibility | 19/30 | 2/14 | n.a. |
| Biochemical findings |  |  |  |
| Reduced GAG synthesis | 9/9 | 4/4<br>(3/4; < HS but > CS/DS) | Reduced GAG production |
| Ultrastructural findings |  |  |  |
| Loosely packed collagen fibrils | 1/1 | n.a. | Disturbed collagen fibril organization in bone and loosely packed collagen fibrils in the dermis |

At 20 dpf, the average weight (W), standard length (SL) and Fulton’s condition factor (*K*), which is used as a measure for the relative proportion of SL to W [29], were not significantly different between *b3galt6*^*−/−*^ and WT siblings (Fig. 3A), although for SL there was a trend towards smaller mutants (P=0.0640). Several externally visible morphological anomalies could be observed in 20 dpf *b3galt6*^*−/−*^ zebrafish as compared to their WT siblings, including a shorter, more compact head, a shorter and rounder frontal region, a shorter parietal/occipital region and a malformed posterior edge of the opercular apparatus (Fig. 3B). Alcian blue (AB) cartilage staining showed no morphological abnormalities, but developmental delay of the head and pectoral and caudal fin endoskeleton was noted (Fig. 3B, Table S7), with cartilage elements being absent or in early development in *b3galt6* KO, while these structures were already fully developed in WT sibling zebrafish. Alizarin red (AR) mineral staining showed delayed mineralization in all regions of the head, the pectoral fin girdle and the caudal fin (Fig. 3B, Table S8), further confirming developmental retardation. In addition, malformations of different mineralized bones, such as the parasphenoid, branchiostegal rays, supracleithrum and the subopercle bone could be detected in the head of *b3galt6*^*−/−*^ zebrafish larvae (Fig. S9). The vertebral column, i.e. the vertebral centra and their associated elements (neural and haemal arches), did not show delayed mineralization in *b3galt6*^*−/−*^ zebrafish larvae, but malformation of the associated elements and fusions of preural vertebrae were observed at a much higher frequency in the *b3galt6*^−/−^ mutants. Kyphosis, scoliosis and lordosis were also more frequently observed and pronounced in *b3galt6*^−/−^ larvae compared to WT siblings (Fig. S10 & Table S8). A similar phenotype was observed in the second *b3galt6* KO model (cmg22 allele), confirming the specificity of the displayed phenotype (Fig. S11).

**Fig. 3.**
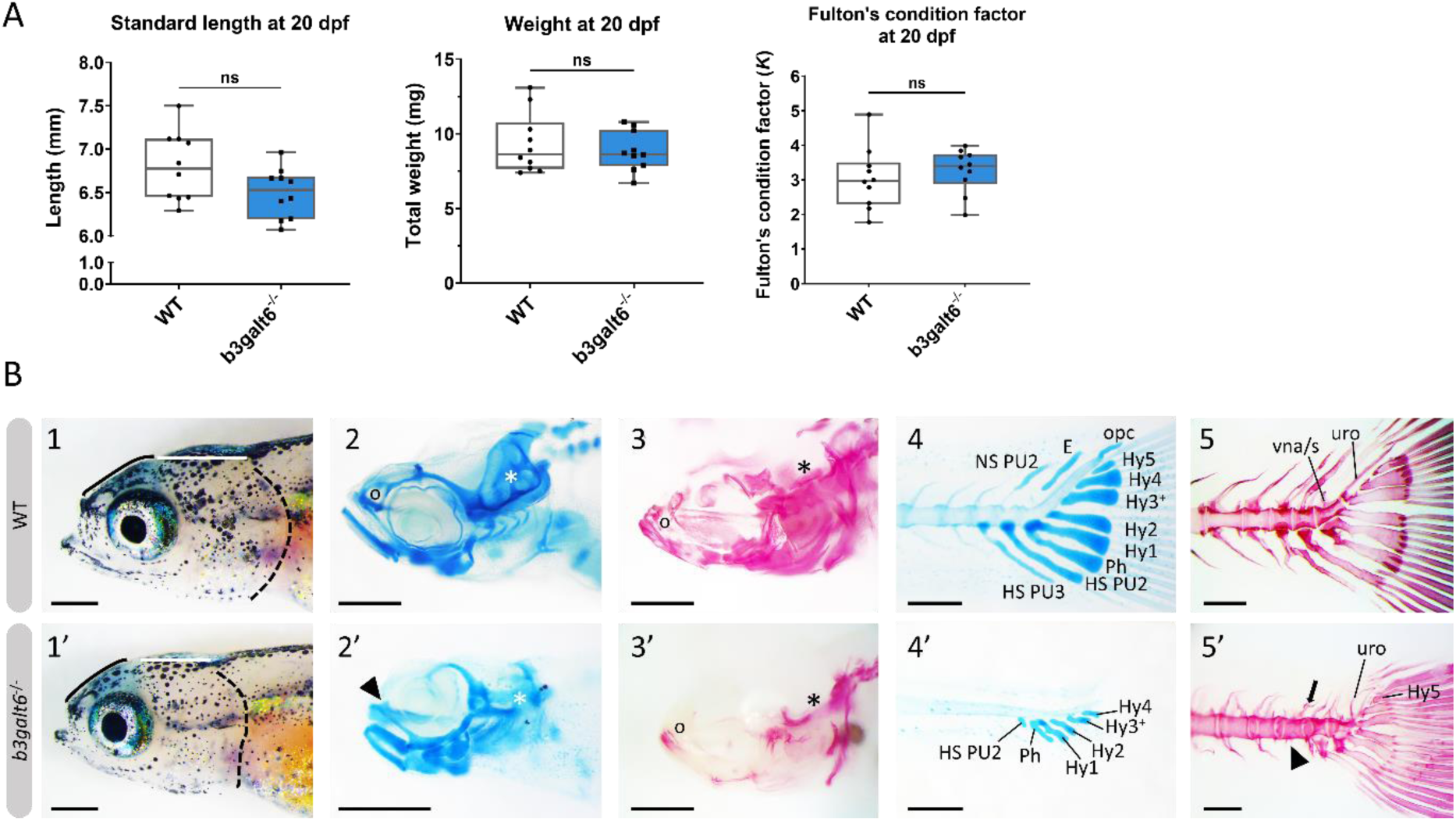
Developmental delay and morphological abnormalities in 20 dpf *b3galt6*^−/−^ zebrafish. (**A**) Weight, standard length and Fulton’s condition factor of WT and *b3galt6*^−/−^ zebrafish larvae at 20 dpf (n=10). (**B**) External phenotype of the head of juvenile WT and *b3galt6*^−/−^ sibling zebrafish. All images are oriented anterior to the left, posterior to the right, dorsal to the top and ventral to the bottom. (**1-1’**) *b3galt6*^−/−^ zebrafish display a rounder frontal region (black line), shorter parietal/occipital region (white horizontal line) and a malformed posterior edge of the opercular apparatus (dotted black line), compared to WT control. (**2-2’**) Representative images of cartilage AB staining of the head. The olfactory region (indicated by ‘o’ in the WT and arrowhead in the *b3galt6*^−/−^ picture) and the otic region (indicated by white asterisk) show delayed development of cartilage elements in *b3galt6*^−/−^ zebrafish as compared to WT. (**3-3’**) Representative images of AR mineral staining of the head. The olfactory region (indicated by ‘o’) and the otic region (indicated by asterisk) show severely delayed mineralization in *b3galt6*^−/−^ zebrafish larvae as compared to WT. (**4-4’**) Representative images of AB staining of the caudal fin. The modified associated elements show developmental delay in *b3galt6*^−/−^ zebrafish as compared to WT. Abbreviations: E, epural; HS PU2, haemal spine of preural 2; HS PU3, haemal spine of preural 3; Hy1-5, hypural 1-5; NS PU2, neural spine preural 2; opc, opistural cartilage; Ph, parhypural (**5-5’**) Representative images of AR mineral staining of the caudal fin. Caudal vertebrae and their associated elements show no delay in ossification. Malformation or absence of caudal fin associated elements (block arrow indicates a neural arch; arrowhead indicates a missing haemal arch), and malformation of uroneural (uro) and vestigial neural arches and spines (vna/s) are observed in *b3galt6*^−/−^ zebrafish; hypurals have a bent appearance. Scale bars: images of the head 500 µm, images of the caudal fin 200 µm.

At the adult stage (four months), *b3galt6*^**−/−**^ zebrafish had a significantly reduced average SL (1.834 ± 0.077 cm), and W (0.1196 ± 0.014 g), when compared to their WT siblings (2.371 ± 0.049 cm, and 0.2497 ± 0.016 g, respectively), while the Fulton’s condition factor (*K*) indicated that SL and W were normally proportioned to each other (Fig. 4A). A similar phenotype was observed in the second *b3galt6* KO model (cmg22 allele), confirming the specificity of the displayed phenotype (Fig. S1).

**Fig. 4.**
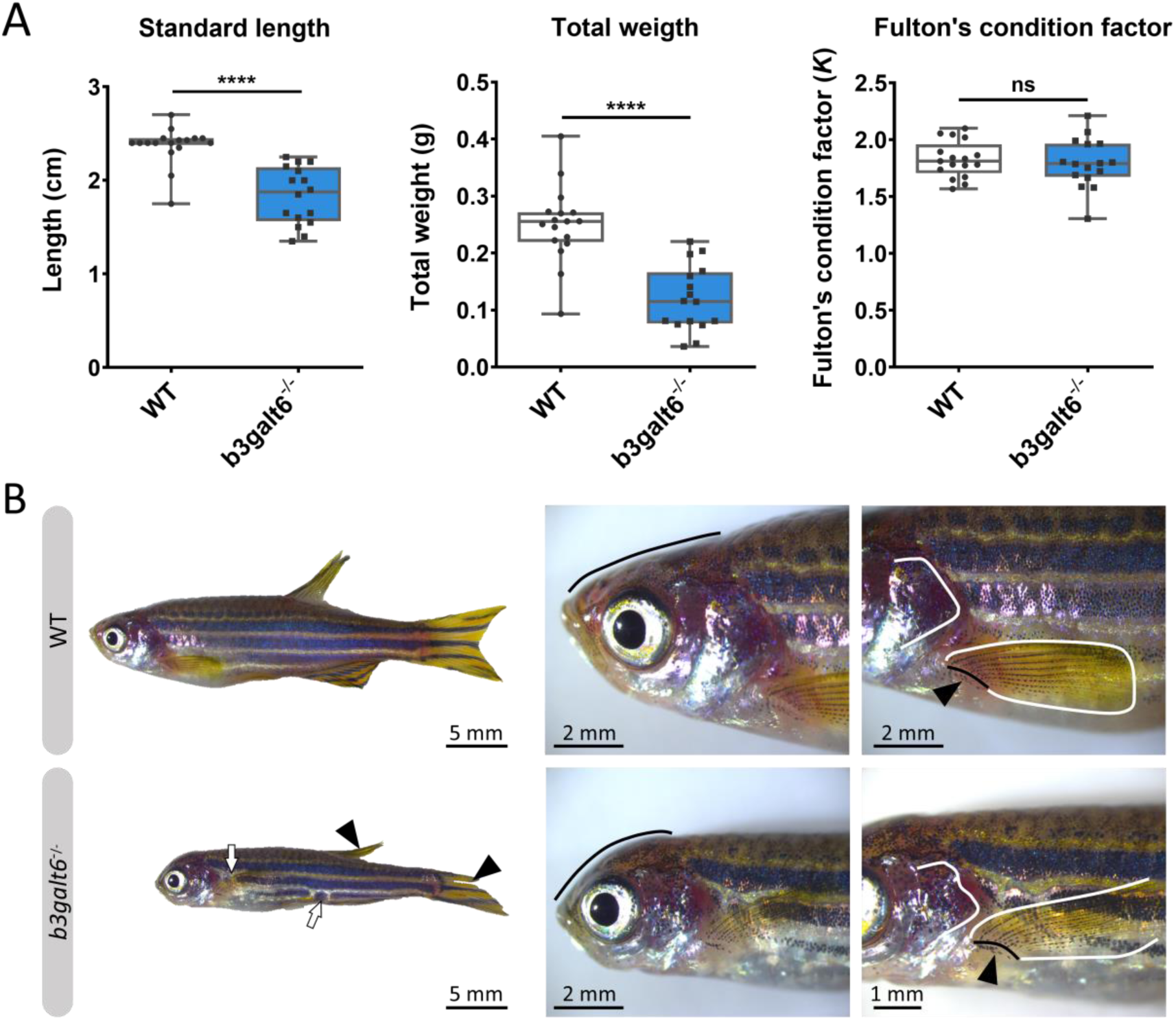
Characterization of the external phenotype of adult *b3galt6*^−/−^ zebrafish. (**A**) *b3galt6*^−/−^ zebrafish (n=16) had a significantly shorter standard length (SL) compared to their WT siblings (n=17). The weight (W) was significantly lower in the mutant zebrafish. The Fulton’s condition factor (*K*= 100*(W/SL^3^)) was not significantly different between *b3galt6*^−/−^ zebrafish and WT siblings. Data are expressed as box plots with min-to-max whiskers on which each individual data point is plotted. An unpaired t-test with Welch’s correction was used to determine significance. ****P<0.0001, ns: not significant. (**B**) Adult *b3galt6*^−/−^ zebrafish exhibit morphological abnormalities compared to their WT siblings; *b3galt6*^−/−^ zebrafish display interruptions of the blue horizontal stripes (white arrows), smaller fins (arrowhead), aberrations of the head shape (black line), of the shape of the opercular apparatus (white line indicates the posterior edge) and of the pectoral fin (black line and arrow indicate the fin rays at the base of the fin, white line shows the outline of the fin).

The head malformations, which were already observed in 20 dpf *b3galt6*^−/−^zebrafish, showed further progression, with a more pronounced smaller, rounded head and an abnormal compaction of the opercular apparatus (Fig. 4B). In addition, a reduced surface area of the fins, with a crenelated postero-distal fin edge was observed in the mutant animals. The pectoral fins were pointing in an abnormal upwards angle (Fig. 4B). Kyphoscoliosis was variably present and mainly occurred in the posterior part of the body (not shown). In addition, an altered pigmentation pattern was observed, mainly on the ventral part of the body where the dark stripes showed interruptions (Fig. 4B).

Further progression of the larval phenotype was also confirmed by AR staining of the mineralized skeleton in adult (four months) *b3galt6*^−/−^ zebrafish. The olfactory region was underdeveloped (Fig. 5A), the cranial roof bones and sutures were deformed (Fig. S12) and the opercular apparatus was smaller and compacted in KO zebrafish (Fig. 5A and Fig. S12). The fifth ceratobranchial arch and its attached teeth were smaller in *b3galt6*^−/−^ zebrafish (Fig. 5B). In the pectoral fins of *b3galt6*^−/−^ zebrafish the radials were smaller, had an irregular shape, and were often fused (Fig. 5B). The autocentra, i.e. the inner layer of the vertebral centra in the vertebral column and caudal fin [30] of KO zebrafish had a normal hourglass shape, but the associated elements often showed presence of extra intramembranous bone (Fig. 5 and Fig. S12) and extra bony elements (Fig. 5A). Fusions were observed in preural vertebrae (Fig. 5A). Lordosis, kyphosis and scoliosis was variably observed along the vertebral column of *b3galt6*^−/−^ mutant zebrafish (not shown). Extra mineralization was observed at the edge (growth zone) of *b3galt6*^−/−^ scales (Fig. 5B). Notably, variation in the severity of the skeletal malformations in *b3galt6*^−/−^ zebrafish was observed.

**Fig. 5.**
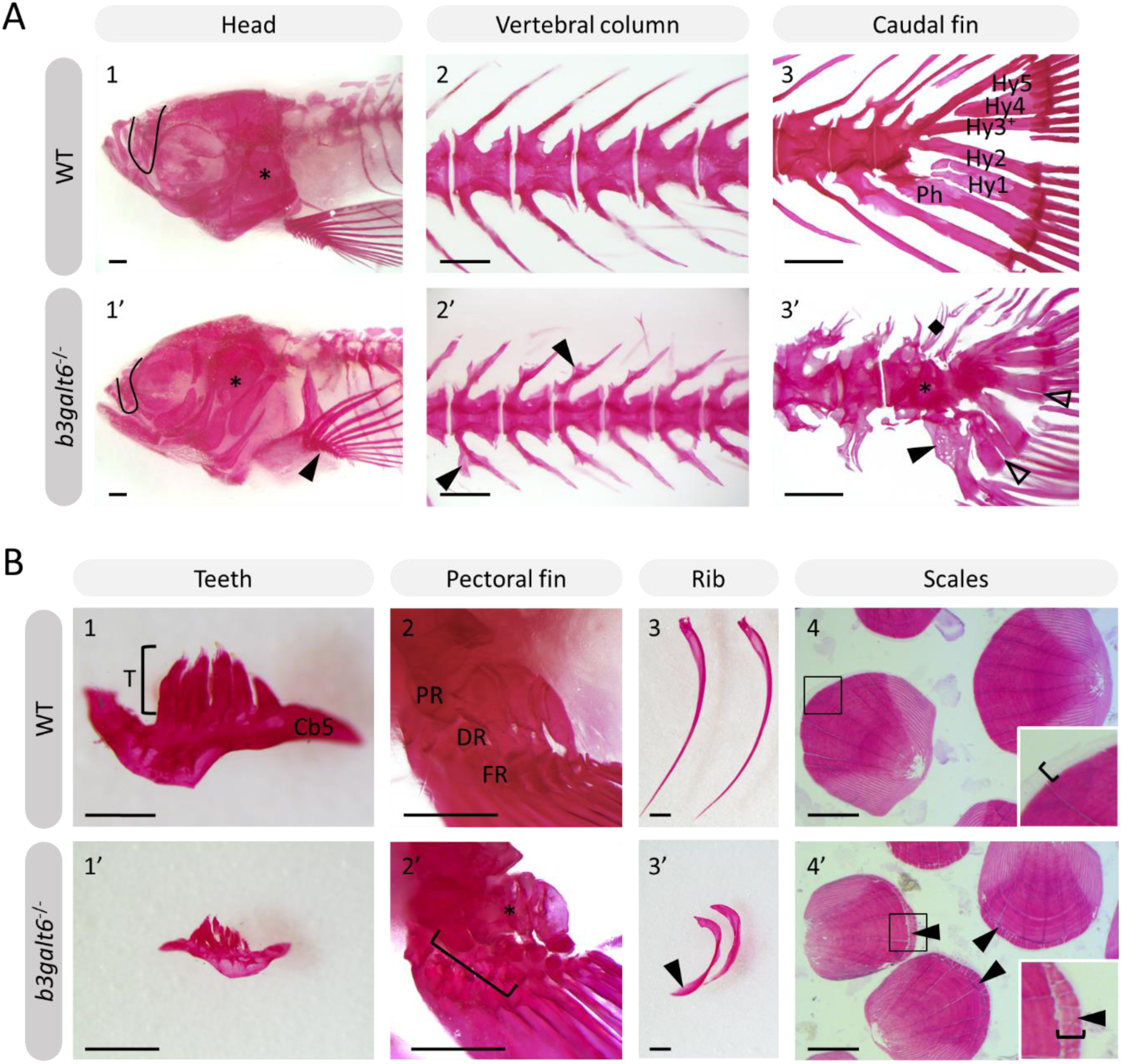
Alizarin red stained mineralized bone in WT and *b3galt6*^−/−^ adult zebrafish. The mineralized bones in the skeleton were studied in four-month-old WT and mutant zebrafish after AR mineral staining. (**A**) *b3galt6*^−/−^ zebrafish display malformation of the cranium, vertebral column and caudal fin. (**A1-1’**) The cranium shows an underdeveloped olfactory region (black line) and a smaller compacted opercular bone (asterisk). (**A2-2’**) The vertebral column shows extra intramembranous outgrowths located where the distal tip of the neural and haemal arches fuse to neural and haemal spine respectively (arrowheads). (**A3-3’**) In the caudal fin fusion of preural vertebrae is observed (asterisk). Associated elements show intramembranous outgrowths (arrowhead) and the presence of extra bony elements (diamond). The distal tip of the hypurals is sometimes split (transparent arrowhead). Structures indicated in the WT caudal fin: Hy1-5, hypural 1-5, Ph, parhypural. (**B**) Details of mineralized structures in *b3galt6*^−/−^ zebrafish. (**B1-1’**) The fifth ceratobranchial arch and its teeth are smaller in KO zebrafish. (**B2-2’**) The proximal (PR) and distal radials (DR) have irregular shapes in mutant zebrafish. Proximal (asterisk) and distal radials are sometimes fused. The bracket indicates a fusion between a proximal and distal radial, and the first fin ray (FR). (**B3-3’**) Ribs also show extra intramembranous bone (black arrowhead). (**B4-4’**) More mineralization was observed at the growth zone (bracket) of the scales in mutant zebrafish (arrowhead). Scale bars = 500 µm.

Subsequent quantitative µCT analysis of four-month-old zebrafish revealed multifaceted effects of B3galt6-deficiency on vertebral morphology and mineralization. Most notably, the volume of vertebral centrum, haemal and neural associated elements was significantly reduced throughout the entire vertebral column in *b3galt6*^−/−^ zebrafish (P<0.01), with more pronounced differences in the abdominal region (Fig. 6). Tissue mineral density (TMD) was not significantly different in the *b3galt6*^−/−^ vertebral bodies compared to WT siblings. However, TMD in zebrafish is proportionally related to standard length [31, 32], and *b3galt6*^−/−^ zebrafish exhibited a significantly reduced body size compared to the WT siblings. We therefore normalized the µCT data to standard length using the normalization procedure outlined by Hur *et al*. [33]. In this procedure, phenotypic data in WT sibling controls was scaled using the mean standard length of *b3galt6*^−/−^ zebrafish as the reference length. Remarkably, after normalization, a significant increase in TMD of centrum, haemal and neural associated elements was noted in *b3galt6*^−/−^ zebrafish compared to WT controls (P<0.001, P<0.01 and P<0.01, respectively) (Fig. S13).

**Fig. 6.**
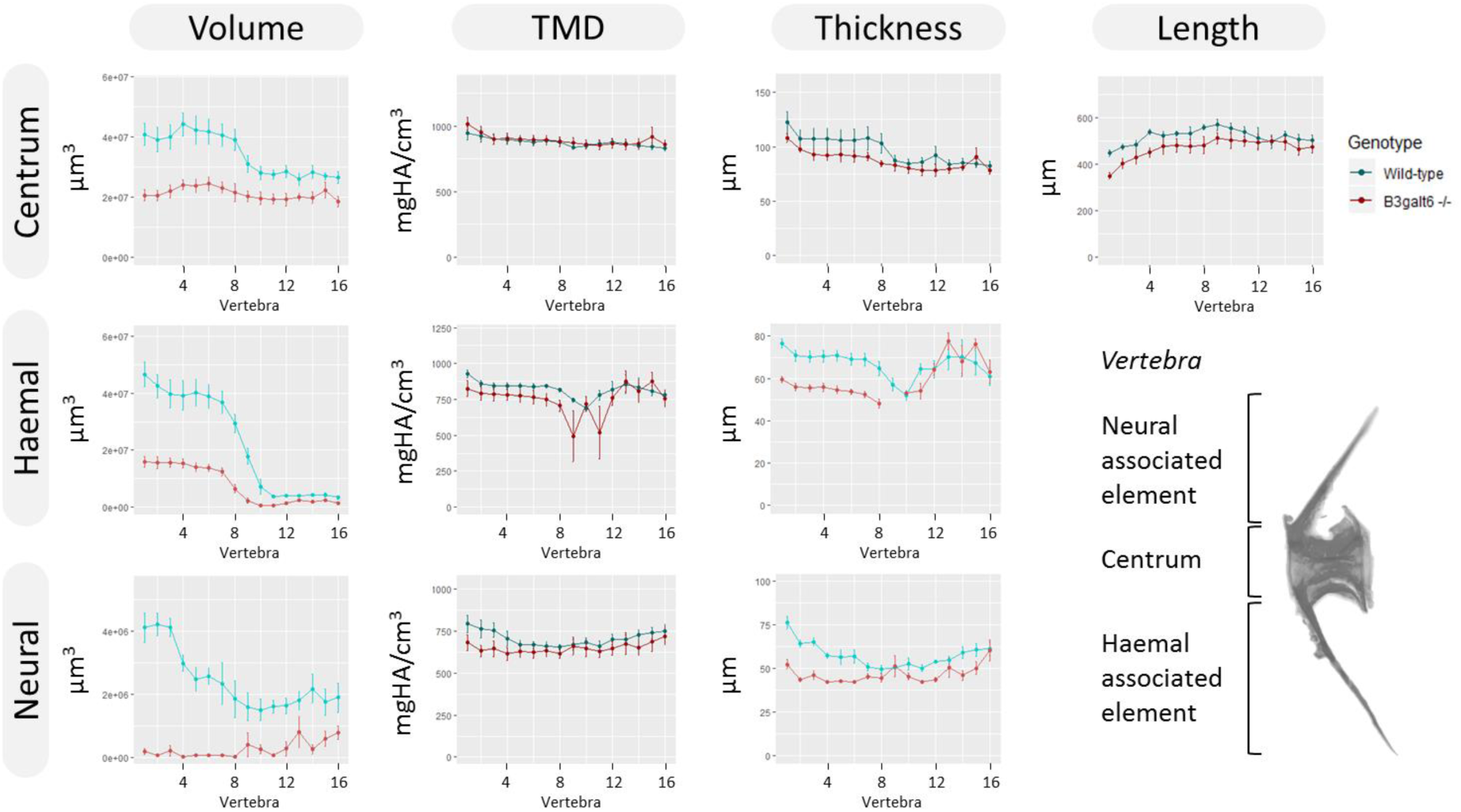
µCT analysis of adult WT and *b3galt6*^−/−^ zebrafish. Quantitative µCT analysis of the vertebral column in five mutant versus five WT zebrafish at the age of four months. The bone volume, tissue mineral density (TMD) and bone thickness were calculated for three parts of each vertebra: the vertebral centrum, the neural associated element (neural arch and neural spine) and the haemal associated element (haemal arch and haemal spine). Plots associated with a significant difference are colored in a lighter coloring scheme. The X-axis represents each vertebra along the anterior-posterior axis. The volume is significantly reduced for each part of the vertebra in *b3galt6*^−/−^ zebrafish (P<0.01, P<0.001 and P<0.000001, respectively). The thickness of the haemal (P<0.01) and neural (P=0.001) associated element is also significantly reduced in the mutants whereas the centrum thickness is not significantly reduced in *b3galt6*^−/−^ zebrafish (P=0.12). TMD is not significantly different between *b3galt6*^−/−^ zebrafish and WT siblings when not normalized for standard length (P>0.2). Data are presented as mean ± SEM. TMD, tissue mineral density; HA, hydroxyapatite.

### *b3galt6* knock-out zebrafish display a disturbed organization of type I collagen fibrils in skin, bone and intervertebral space

Transmission electron microscopy (TEM) studies on the dermis of spEDS-*B3GALT6* patients revealed an abnormal collagen fibril architecture with loosely packed collagen fibrils of variable size and shape [5]. We therefore studied the effect of loss of galactosyltransferase II activity on collagen fibril architecture. TEM sections of skin biopsies of *b3galt6*^−/−^ zebrafish showed an abnormal dermal collagen fibril architecture characterized by loosely packed collagen fibrils and larger and more interfibrillar spaces compared to WT siblings (Fig. 7A). In addition, at the level of the scales a thicker epidermal layer was observed and electron dense structures were present in the interfibrillar spaces (Fig. 7A). Next, we examined the intervertebral region and adjacent vertebral centra in the vertebral column of five-month-old zebrafish by means of TEM. In general, the type I collagen fibers in *b3galt6*^-*/-*^ zebrafish displayed a lower degree of organization (Fig. 7B). This was observed in immature collagen layers, deposited by osteoblasts in the outer edges of the intervertebral space, but also in the mature type I collagen layers of the intervertebral ligament and in the autocentrum, the inner layer of the vertebral bone, where a lack of the typical plywood-like organization was seen. Detailed investigation of TEM sections of the vertebral bone revealed presence of several electron dense spots of unknown origin in knock-outs. These ‘dark spots’ may reflect extrafibrillar hydroxyapatite crystals (Fig. 7B).

**Fig. 7.**
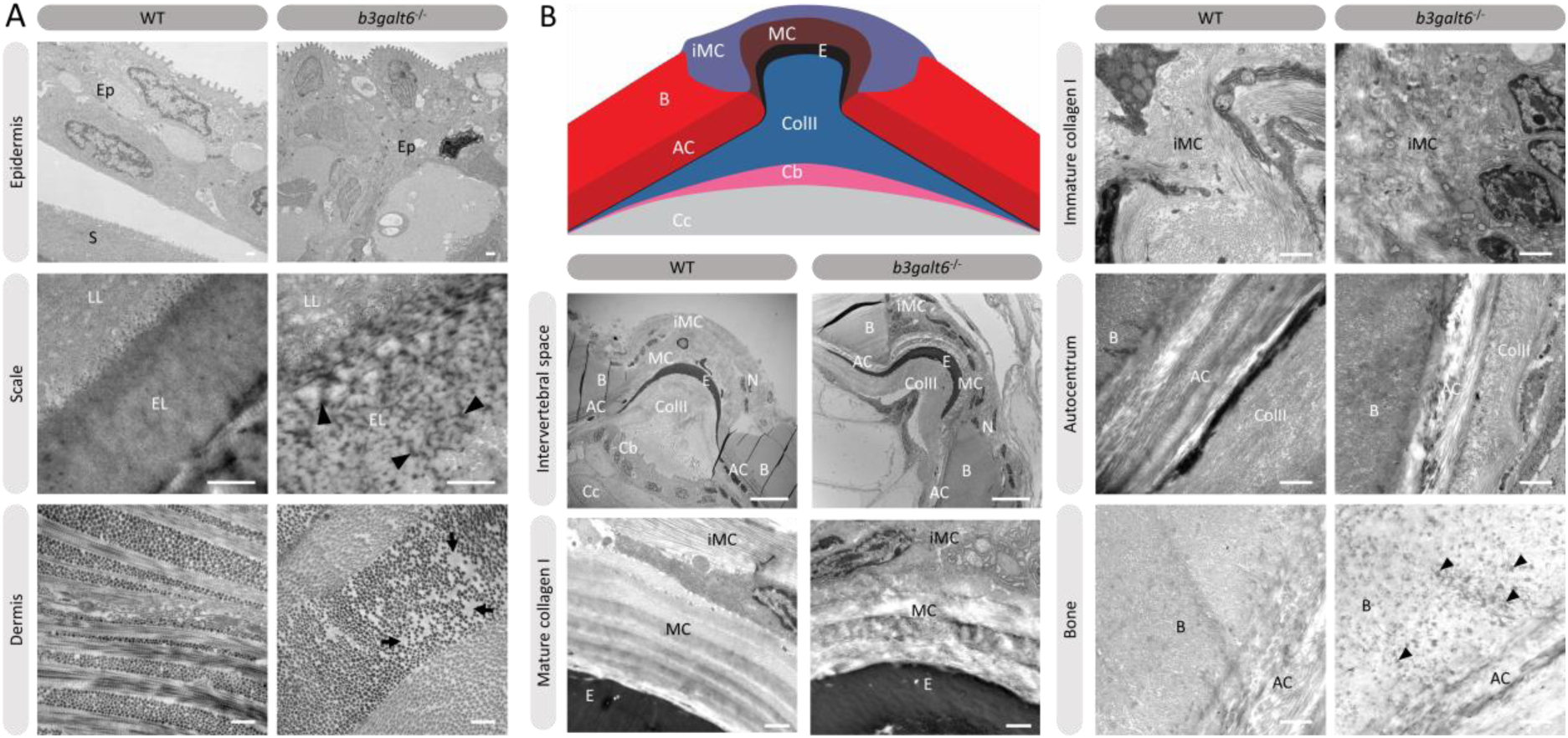
Collagen fibrillar architecture in five-month-old adult zebrafish. (**A**) Skin from five-month-old WT and *b3galt6*^−/−^ adult zebrafish. The epidermis (Ep), the outer layer of the skin covering the scales (S) of zebrafish, appears thicker in *b3galt6*^−/−^ zebrafish compared to the epidermis of WT zebrafish. A mature zebrafish scale consists of several layers. The limiting layer of the scale (LL) is rich in fine granules that in some areas are linearly arranged, forming dense layers. This matrix contrasts with that of the external layer(s) (EL) below. Structures resembling extrafibrillar crystals (arrowhead) are noted in the external layer of the mutant scale, which are rare in WT zebrafish sections. Upon investigation of the dermis of *b3galt6*^−/−^ mutants, increased interfibrillar spaces are noted (arrows), which are absent in the dermis of WT adults. Scale bars = 500 nm. (**B**) Schematic representation and TEM images of the zebrafish intervertebral space. Schematic representation of the zebrafish intervertebral space. Abbreviations: AC, autocentrum; B, bone; Cb, chordoblasts; Cc, chordocytes; ColII, type II collagen; E, elastin; iMC, immature collagen type I layer; MC, mature collagen type I layer. Starting from the central body axis, several recognizable structures are present in the intervertebral space along a proximo-distal axis: (i) notochord tissue consisting of chordocytes and chordoblasts, (ii) the notochord sheath consisting of a type II collagen layer and an external elastic membrane, (iii) and a connective tissue ligament consistent of a mature and an immature type I collagen layer. The mature type I collagen layer is organized in a typical plywood-like pattern and is continuous with the inner layer of the vertebral bone, i.e. the autocentrum, which also shows a plywood-like organization. The immature type I collagen layers consist of loose collagen fibers connecting the outer layer of vertebral bones. Notice the higher number of nuclei (N) in the *b3galt6*^−/−^ zebrafish at the intervertebral space. Mature collagen (MC) displays a typical plywood-like organization, in WT zebrafish but not in comparable regions for *b3galt6*^−/−^ mutant zebrafish. Cross section and longitudinal type I collagen regions are less clearly demarcated and smaller in immature collagen (iMC) from *b3galt6*^−/−^ zebrafish compared to WT siblings. It is more difficult to pinpoint the different layers of the autocentrum (AC) in the KO compared to WT zebrafish. Bone (B) from the *b3galt6*^−/−^ zebrafish shows electron dense structures (arrowhead). Scale bars = 10 µm for the intervertebral space and 1 µm for the other images.

Mass spectrometry analysis of type I collagen modification in bone derived from the vertebral column showed no significant difference in hydroxylation status of the α1(I) K87 and C-telopeptide lysine residues (Table S9). These lysine residues are conserved in zebrafish and human type I collagen and are crucial for intermolecular crosslinking of collagen fibrils in bone matrix [34].

### *b3galt6* knock-out zebrafish display ultrastructural muscle abnormalities and lower endurance

Human spEDS-*B3GALT6* patients often suffer from muscle hypotonia with a delay in motor development. In view of the observed reduction in the concentration of the different GAG moieties in muscle of *b3galt6*^−/−^ zebrafish, we investigated the function and the ultrastructural appearance of muscle.

First, the critical swimming speed (*U*_crit_) and endurance of the zebrafish were assessed by means of a custom-made swim tunnel. The *U*_crit_, defined as the maximum velocity that the tested zebrafish could sustain for at least 5 minutes [35], was significantly lower in *b3galt6*^−/−^ zebrafish (30.1 ± 2.39 cm/s) compared to length-matched WT (41.61 ± 1.942cm/s) (P<0.01) (Fig. 8A). The stamina of each zebrafish was determined by an effort test where it was tested how long zebrafish could swim at high speed (37 cm/s) after a three-minute warm-up. This effort test showed that the *b3galt6*^−/−^ zebrafish performed less well compared to length-matched WT zebrafish, indicating a lower endurance (Fig. 8B).

**Fig. 8.**
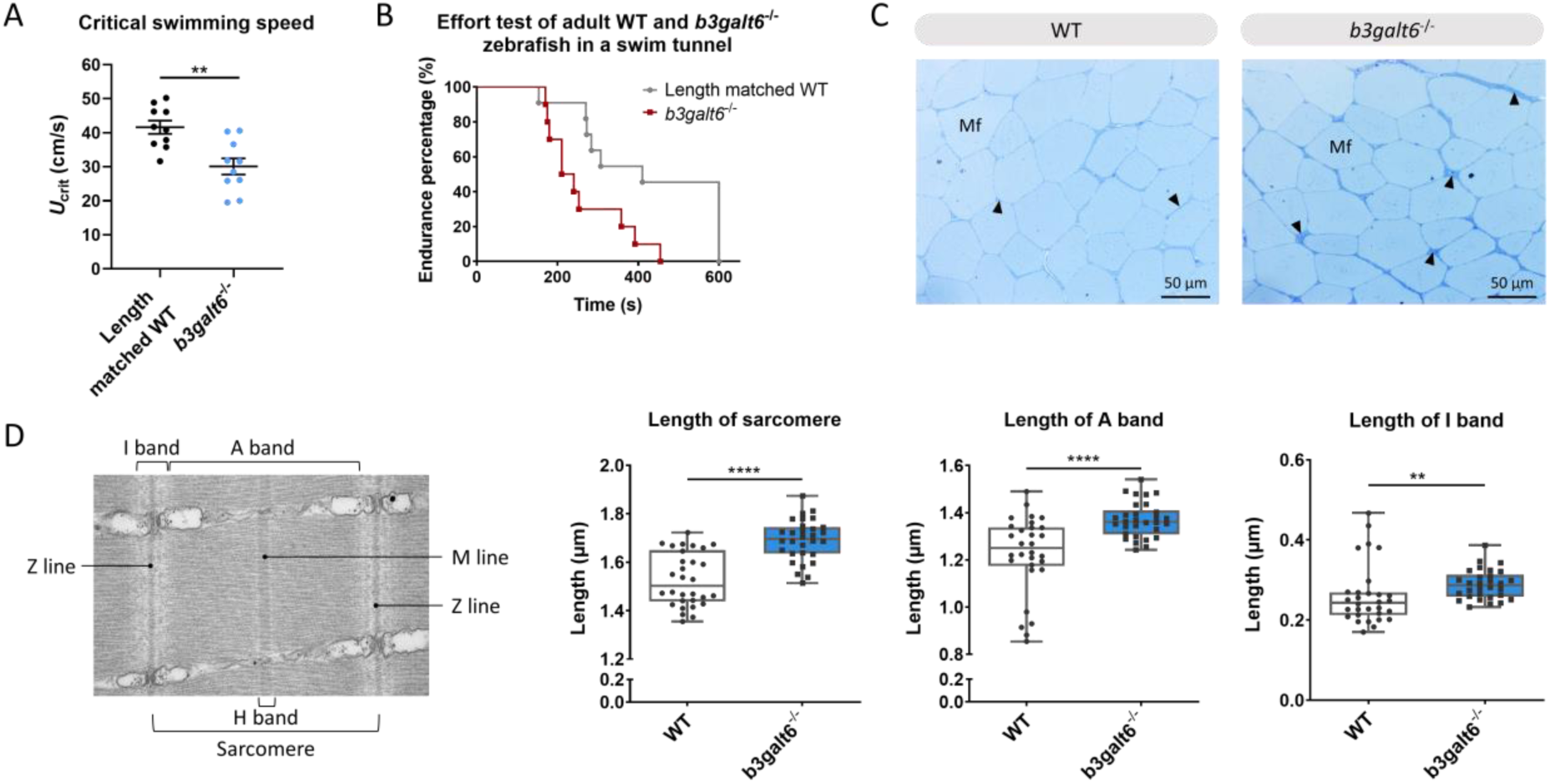
Functional and structural analysis of adult WT and *b3galt6*^−/−^ zebrafish muscle. (**A**) The critical swimming speed and (**B**) the endurance performance was determined for *b3galt6*^−/−^ and length-matched WT zebrafish. (**C**) Semi-thin cross sections, stained with toluidine blue, demonstrate the presence of a thicker endomysium (arrowhead), located between the muscle fibers (Mf) at the horizontal myoseptum of *b3galt6*^−/−^ zebrafish at the age of five months. (**D**) Transmission electron microscope structure of a sarcomere with indications of the Z and M line and A, I and H band. The sarcomere length was determined at 5 different loci per semi-thin section, for 6 semi-thin originating coming from 3 different zebrafish per genotype (N=30). A significant increase in sarcomere length for *b3galt6*^−/−^ is noted. Furthermore, both the A and I band show an increase in length for the mutant zebrafish. Data are expressed as box plots with min-to-max whiskers on which each individual data point is plotted. A Mann-Whitney U test was used to determine significance. ****P<0.0001, **P<0.01.

The muscle tissue in the zebrafish trunk was further investigated at the ultrastructural level. Toluidine blue-stained semi-thin cross sections revealed that the endomysium, a layer of connective tissue ensheating the muscle fibers, was enlarged along the horizontal myoseptum of *b3galt6*^−/−^ zebrafish, although no clear differences in muscle fiber diameter were noted (Fig. 8C). Ultra-thin parasaggital sections, investigated by TEM, showed that the length of the sarcomeres, the smallest basic units of striated muscle, as well as the length of the A and I band were significantly increased in the *b3galt6*^−/−^ zebrafish (Fig. 8D).

## Discussion

In this study we developed two viable *b3galt6* KO zebrafish models using CRISPR/Cas9 mutagenesis, and performed an in-depth characterization of the biochemical, ultrastructural and phenotypical consequences of β3GalT6-deficiency on different tissues, including skin, cartilage, mineralized bone and muscle. These models currently represent the only vertebrate models available for the study of the pathogenetic consequences of β3GalT6-deficiency on tissue development and homeostasis. In both models we detected a significant reduction, yet not a complete absence in *b3galt6* mRNA expression (Fig. 1E). One possible explanation is the absence of canonical nonsense-mediated decay (NMD) in the single-exon *b3galt6* gene, due to a lack of exon junction complexes (EJCs) [36]. We were unable to examine the effect of the mutations on B3galt6 protein levels and localization due to the unavailability of a suitable antibody. Nevertheless, since the cmg20 and cmg22 alleles are predicted to introduce a PTC respectively upstream and in the catalytic domain, residual translation of some mRNA into a truncated protein is not expected to lead to the production of an active enzyme. β3galt6 is a key enzyme in the biosynthesis of the common linkage region of PGs, and reduced levels of active enzyme are thus expected to equally affect the biosynthesis of HS and CS/DS GAG chains. We therefore quantified the disaccharide concentrations in bone, muscle and skin of adult *b3galt6*^−/−^ zebrafish by means of a combination of enzymatic digestion and anion-exchange HPLC. This revealed a distinct, and relatively proportional decrease in the concentration of CS, DS, hybrid CS/DS and HS moieties in all examined tissues, compared to age-matched WT zebrafish (Table 1). Nonetheless, low amounts of GAGs were still produced in the KO zebrafish in absence of upregulation of other galactosyltransferase member(s) from the *b3galt* family or by other linkage-enzymes (Fig. S6). Together with the previous demonstration that the other five β3GalT human enzymes (e.g. β3GalT1, β3GalT2, β3GalT3/β3GalNAcT1, β3GalT4 and β3GalT5) are involved in the biosynthesis of glycoproteins and glycolipids [11], compensation for galactosyltransferase II deficiency in zebrafish by other *b3galt* family members is unlikely. Chinese hamster ovary *B3galt6* KO cells were also shown to be capable of producing GAGs, in contrast to *Xylt2, B4galt7* or *B3gat3* KO cells [37]. The recent identification of the presence of a non-canonical trisaccharide linkage region, lacking one Gal residue (GlcA-Gal-Xyl-*O*-), as a minor constituent in the PG bikunin in the urine of healthy human individuals [28], prompted us to examine the composition of the GAG linkage region in our *b3galt6* KO zebrafish models, by analyzing protein extracts from adult *b3galt6*^−/−^ and WT zebrafish via LC-MS/MS. This led to the striking finding that the few detected PGs in the *b3galt6*^−/−^ zebrafish all presented with a trisaccharide PG linkage region, consisting of only one Gal (Fig. 2B) (Table S6). Further *in vitro* analysis indicated that GlcAT-I/B3GAT3 is capable of adding a GlcA to Gal-Xyl-*O*-(Fig. S8). This observation might explain the phenotypic differences observed between GalT-II/B3GALT6 deficiency and deficiencies of the other linkage region enzymes, for which such a rescue mechanism might be absent. This hypothesis is supported by the lack of GAGs in the Chinese hamster ovary *Xylt2, B4galt7* or *B3gat3* KO cells and by the lethal phenotype observed in the *b4galt7* KO zebrafish model [38]. At present, we have no information on the length of the GAG chains attached to the trisaccharide linkage region or the composition and the possible modifications of these HS and CS/DS chains. As such, the GAG structure, length and modifications in the *b3galt6*^−/−^ zebrafish might be different from the ones in WT zebrafish.

HSPGs and CS/DSPGs are essential macromolecules for the development, signaling and homeostasis of many tissues. It therefore comes as no surprise that defects in linkage region biosynthesis can affect many organs. Endochondral ossification of cartilage (forming chondral bones) for example is a multi-staged process that must be coordinated in space and time. CS/DSPGs and HSPGS, which have key roles in modulating signaling events during vertebrate skeletal development, are expressed in a spatially and temporally highly controlled manner [24], thus regulating the timing of endochondral ossification. Genetic defects disrupting PG synthesis or structure have been shown to disrupt the timing of skeletal development. Two zebrafish models with abnormal PG linkage region biosynthesis, i.e. the *fam20b* and *xylt1*-deficient models, revealed partial decrease in CSPGs, which are predominantly present in cartilage, and accelerated endochondral ossification, showing that CSPGs can negatively regulate skeletogenic timing [24]. On the other hand, a strong reduction in CSPGs and partial decrease in HSPGs was noted in *b3gat3* mutant zebrafish, but this led to a delay in endochondral ossification and short and disorganized cartilaginous structures [39]. Our B3galt6-deficient zebrafish models also show a delay in skeletal development at juvenile stages. At 20 dpf, a delay was noticed in the development of cartilage and mineralized bony elements of the craniofacial structures, the pectoral and caudal fin endoskeleton (Fig. 3B) (Tables S7 and S8), and these abnormalities had progressed in adult zebrafish. The reduced body length, misshapen cranial bones and smaller teeth observed in the head skeleton, and general skeletal dysplasia with kyphosis and scoliosis seen in the vertebral column of *b3galt6*^−/−^ adult zebrafish is reminiscent of the skeletal phenotype observed in human β3GalT6-deficient individuals (Table 2).

In addition to skeletal dysplasia, we also observed abnormalities in bone mineralization. Adult mutant zebrafish stained with AR showed the presence of extra intramembranous bone and extra bony elements on the associated elements of the vertebral centra. On the other hand, quantitative µCT analysis of the vertebral column and associated elements revealed a generalized reduction in bone volume and thickness, and a relative increase in TMD after correction for reduced body length of the mutant zebrafish (Fig. 6 and Fig. S13). Although it is not clear so far if the increased fracture risk in spEDS-*B3GALT6* patients can be attributed to a lower bone volume combined with a higher TMD, such hypothesis has been put forward for the brittle bone disorder Osteogenesis Imperfecta (OI) based on observations in both patients and animal models [40, 41]. In the Chihuahua (*col1a1a*^chi/+^) OI zebrafish model the intrafibrillar mineral platelets were shown to be smaller and more densely packed, leading to an increased mineral to matrix ratio, and consequently a reduced elastic modulus-to-hardness ratio, a measure for fracture toughness [41-43]. In addition, in both OI patients and the mouse model *oim*, reduced collagen matrix organization was shown to lead to the formation of extrafibrillar crystals, which are larger compared to intrafibrillar crystals [43]. Eventually, additional minerals stabilize the tissue, but they do so at the cost of increased brittleness of the bone [44]. In accordance, in *b3galt6*^−/−^ zebrafish we observed an irregular collagen organization interspersed with large interfibrillar spaces and electron dense spots on ultra-thin sections of skin and bone via TEM. The ‘dark spots’ observed in the vertebral bone could be remnants of extrafibrillar mineral aggregates, contributing to increased mineralization but also increased brittleness of the skeleton.

Abnormalities in collagen fibril organization were observed in many mutant zebrafish tissues. Besides vertebral bone, irregular collagen organization was also observed in the intervertebral space, intervertebral ligament, and the dermis. (Fig. 7). This strongly resembles the previously reported abnormal dermal collagen fibril architecture in the skin from a β3GalT6-deficient individual [5]. This could be attributed to abnormalities in GAG chains of small leucine-rich PGs (SLRPs), such as the DSPGs decorin and biglycan, which are known to be of particular importance for regulating collagen fibril organization in connective tissues through their core protein and GAG chains [45]. SLRPs are also thought to influence collagen cross-linking patterns, but the mechanism by which they do so remain largely unknown [45]. Although similar hydroxylation levels were observed at K87 and the C-telopeptide of the type I collagen α1 chain in *b3galt6*^−/−^ and WT adult zebrafish (Table S9), we cannot rule out that type I collagen cross-linking might still be affected.

In view of the observation that human β3GalT6-deficient individuals often present quite severe muscle hypotonia with delayed gross motor development and reduced endurance, we also studied the effect of B3galt6-deficiency on GAG synthesis in muscle, and on muscle function and ultrastructure (Fig. 8). Similar to the other examined tissues, HS and CS/DS GAG concentrations were severely reduced in muscle tissue. It is known that the ECM in skeletal muscle plays an important role in muscle fiber force transmission, maintenance and repair. Numerous PGs, including the HSPGs syndecans 1-4, glypican-1, perlecan and agrin, and CS/DSPGs, such as decorin and biglycan, are present in muscle ECM. Besides their structural role, many of them are known to bind growth factors (e.g. TGFβ). Those growth factors can be stored and released by the negatively charged GAGs. Certain enzymes in the ECM can cleave GAG chains, thereby releasing associated growth factors, allowing their interaction in cell signaling and mechanotransduction [46]. The significant decrease in critical swimming speed (*U*_crit_) and endurance observed in the *b3galt6*^−/−^ zebrafish suggest decreased muscle function (Fig. 8A-B). Further evidence for an intrinsic muscle dysfunction in the *b3galt6*^−/−^ model comes from semi-thin muscle sections, showing an enlargement of the connective tissue sheet surrounding the muscle fibers (Fig. 8C). Nevertheless, we cannot exclude that the observed skeletal malformations and/or other abnormalities, such as respiratory or neuronal dysfunction, could also influence the outcome of these tests.

Taken together, we generated and characterized the first *b3galt6* KO zebrafish models and show that their phenotype largely mimics that of human β3Galt6-deficiency. A strong reduction in HS, CS and DS GAG concentration was observed in *b3galt6*^−/−^ zebrafish. Yet, we did observe a small amount of GAGs being produced with a unique trisaccharide PG linkage region. Furthermore, we showed that a lack of B3galt6 causes disturbances in collagen fibril organization in skin, ligament and bone, with the latter also showing abnormalities in mineralization, and leads to structural and functional abnormalities of the muscle. These KO zebrafish can serve as useful models to further unravel the underlying pathogenic mechanisms of this complex multisystemic disorder and serve as a tool for the development and evaluation of possible therapeutic interventions, such as the stimulation of intrinsic compensational mechanisms for the production of PGs.

## Material and Methods

### Zebrafish Maintenance

Wild-type (AB strain) and *b3galt6* knock-out zebrafish lines were reared and maintained by standard protocols [48]. All animal studies were performed in agreement with EU Directive 2010/63/EU for animals, permit number: ECD 16/18. All zebrafish used in a single experiment were bred at the same density. All efforts were made to minimize pain, distress and discomfort.

### Alcian blue and alizarin red staining

Twenty-day-old larvae (n=10) and five-month-old WT and *b3galt6*^*−/−*^ (n=5) adult zebrafish were stained as previously described [38]. See Supplementary Information for details.

### µCT scanning analysis

Whole-body µCT scans of four-month-old WT (n=5) and *b3galt6*^−/−^ (n=5) siblings were acquired on a SkyScan 1275 (Bruker, Kontich, Belgium) using the following scan parameters: 0.25 mm aluminum filter, 50 kV, 160 µA, 65 ms integration time, 0.5° rotation step, 721 projections/360°, 5 m 57 s scan duration and 21 µm voxel size. DICOM files of individual zebrafish were segmented in MATLAB using custom FishCuT software and data were analyzed and corrected for standard length in the R statistical environment, as previously described [33].

### Histology and transmission electron microscopy

Zebrafish bone, consisting of the last transitional vertebra and two consecutive caudal vertebrae [49, 50], skin, at the lateral line below the dorsal fin, and muscle samples, located dorsolateral underneath the dorsal fin, dissected from five-month-old WT (n=5) and *b3galt6*^−/−^ (n=5) zebrafish, were fixed and embedded in epon following the procedures outlined by Huysseune and Sire [51].

### Swim tunnel experiments

Ten WT (five months) and ten *b3galt6*^*−/−*^ zebrafish (eight months), were selected based on their similar (not significantly different) length and weight. The critical swimming speed (*U*_*crit*_) was measured in a swim tunnel for both WT and *b3galt6*^−/−^ zebrafish as previously described [52]. Furthermore, the endurance of each zebrafish was tested by means of an effort test in the swim tunnel. Each zebrafish had a three minute warm-up and after the third minute, the speed increased until 37 cm/s for maximum 7 minutes. When the zebrafish was not able to swim anymore, the test was stopped.

### Collagen crosslinking analysis

The collagen crosslinking was analyzed in the dissected vertebral column (excluding Weberian apparatus and urostyle) from five-month-old wild-type (n=5) and *b3galt6*^−/−^ (n=5) siblings as described before in Gistelinck *et al*. [40].

### Disaccharide composition analysis of CS, DS, and HS chains

The level of total disaccharides from CS, DS, and HS in bone, muscle and skin tissues from nine-month-old zebrafish (n=5) were determined as described previously [53, 54].

### Glycopeptide preparation and analysis

#### LC-MS/MS sample preparation

CS- and HS-glycopeptides were purified from six-month-old WT (n=3) and *b3galt6*^−/−^ (n=3) zebrafish using a combination of extraction, trypsin digestion and anion exchange chromatography, slightly modified (see Supplementary Information) from previously described protocols [27, 55].

#### LC-MS/MS analysis

The samples were analyzed on an Orbitrap Fusion mass spectrometer coupled to an Easy-nLC 1200 liquid chromatography system (Thermo Fisher Scientific., Waltham, MA) (see Supplementary Information). Nanospray Flex ion source was operated in positive ionization mode and three separate higher-energy collision-induced dissociation (HCD) MS/MS spectra were recorded for each precursor ion [27, 55].

#### LC-MS/MS data analysis

The glycopeptide files were analyzed using both manual interpretation and automated Mascot searches for GAG-glycopeptides. The files were manually interpreted using the Xcalibur software (Thermo Fisher Scientific). Database searches were performed against *Danio rerio* in the UniProtKB/Swiss-Prot database and NCBI using Mascot Distiller (version 2.6.1.0, Matrix Science, London, U.K) and an in-house Mascot server (version 2.5.1). Glycan structures with very small mass differences were manually evaluated at the MS2-level.

### GlcAT-I assay

GlcA-transferase assay was carried out by the methodology as previously described [12, 56, 57], with slight modifications.

Detailed information regarding the procedures used is provided in SI Appendix, Supplementary Material and Methods.

## Acknowledgements

We would like to thank the TEM facility of the Nematology Research Unit, member of the UGent TEM-Expertise center (life sciences). We are grateful to Hanna De Saffel and Petra Vermassen for their technical assistance on processing the TEM samples. The LC-MS/MS analyses were performed at the BioMS node at the Proteomics Core Facility, Sahlgrenska Academy, University of Gothenburg with support from the Swedish National Infrastructure for Biological Mass Spectrometry (BioMS), funded by the Swedish Research Council. This manuscript has been released as a pre-print at bioRxiv, (Delbaere et al.) [58].

