## Supplemental Information for "*b3galt6* knock-out zebrafish recapitulate β3GalT6-deficiency disorders in human and reveal a trisaccharide proteoglycan linkage region"

### Supplementary information

#### Supplementary Results

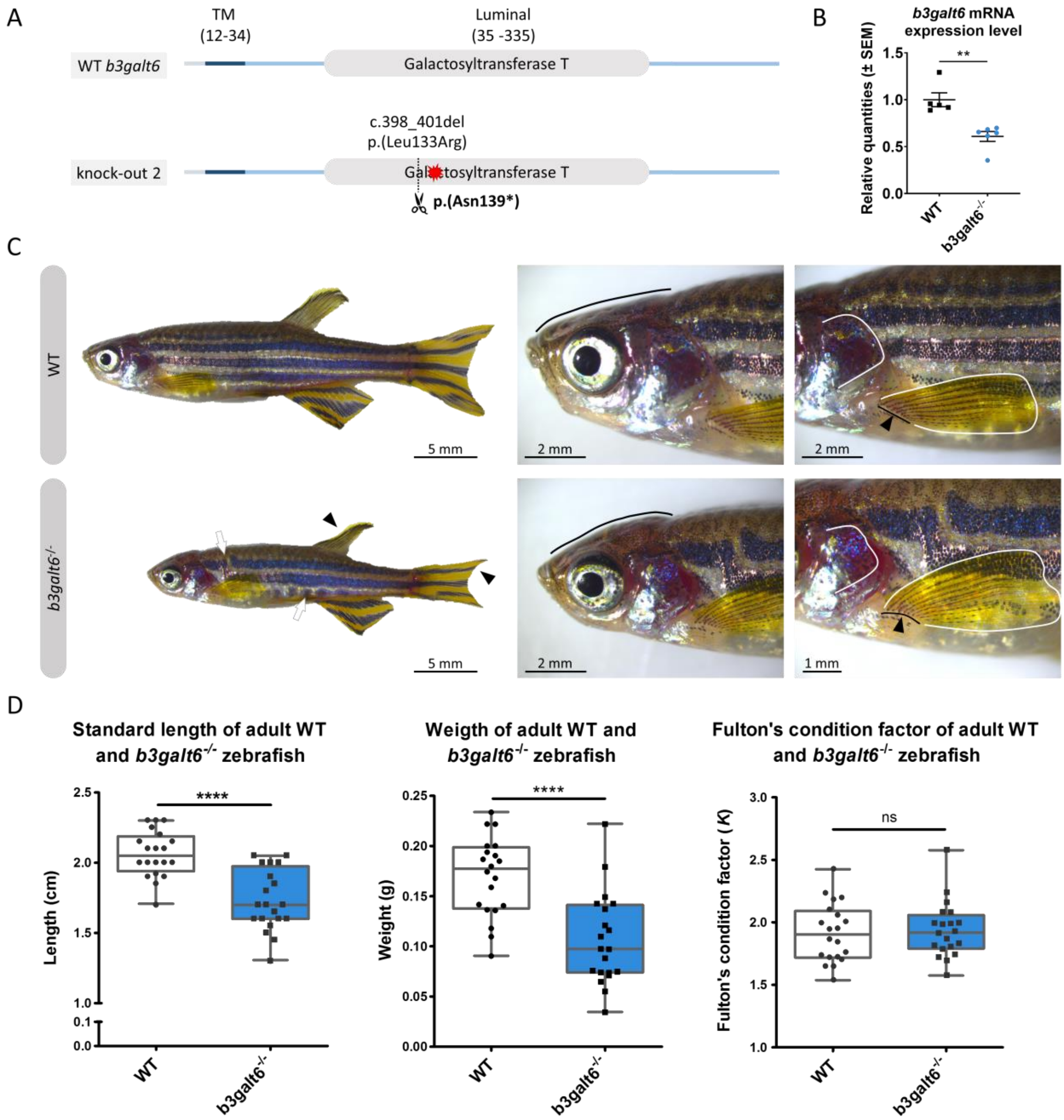


**Fig. S1. Characterization of the second CRISPR/Cas9 generated *b3galt6*$^{-/-}$ zebrafish harboring cmg22 allele.** (**A**) Schematic representation of the frameshift mutation in the second *b3galt6* KO mutant. B3galt6 consists of three parts, a cytosolic domain (amino acid (AA) 1-11), a transmembrane domain

(TM, AA 12-34) and a luminal domain (AA 35-335). A frameshift mutation, c.389_402del (p.(Leu133Arg)), was inserted by the CRISPR/Cas9 gene editing technique, leading to a premature stop codon (p.(Asn139*)) in the catalytic domain (AA 81-268). (**B**) RT-qPCR analysis of relative *b3galt6* mRNA expression levels in *b3galt6*$^{-/-}$ adult zebrafish (0.6087 ± 0.05238) (n=6) compared to WT siblings (1 ± 0.07387) (n=5). Data are expressed as mean ± SEM. An unpaired t-test was used to determine significance, **P<0.01. (**C**) Adult *b3galt6*$^{-/-}$ zebrafish exhibit abnormalities of the external phenotype compared to their wild-type siblings. *b3galt6*$^{-/-}$ zebrafish display aberrations that are most apparent in the head (black line), the jaws, the opercular apparatus (white line), fins (black arrowheads) and pigmentation pattern (white arrow). (**D**) *b3galt6*$^{-/-}$ zebrafish had a significantly shorter standard length (SL) compared to their WT siblings. The weight (W) was strongly diminished in the mutant zebrafish. The Fulton's condition factor ($K$= 100*(W/SL$^3$)) was not significantly different between *b3galt6*$^{-/-}$ zebrafish and WT siblings (P=0.7379). Data are expressed as box plots with min-to-max whiskers on which each individual data point is plotted. An unpaired t-test with Welch's correction was used to determine significance. ****P<0.0001, ns: not significant.

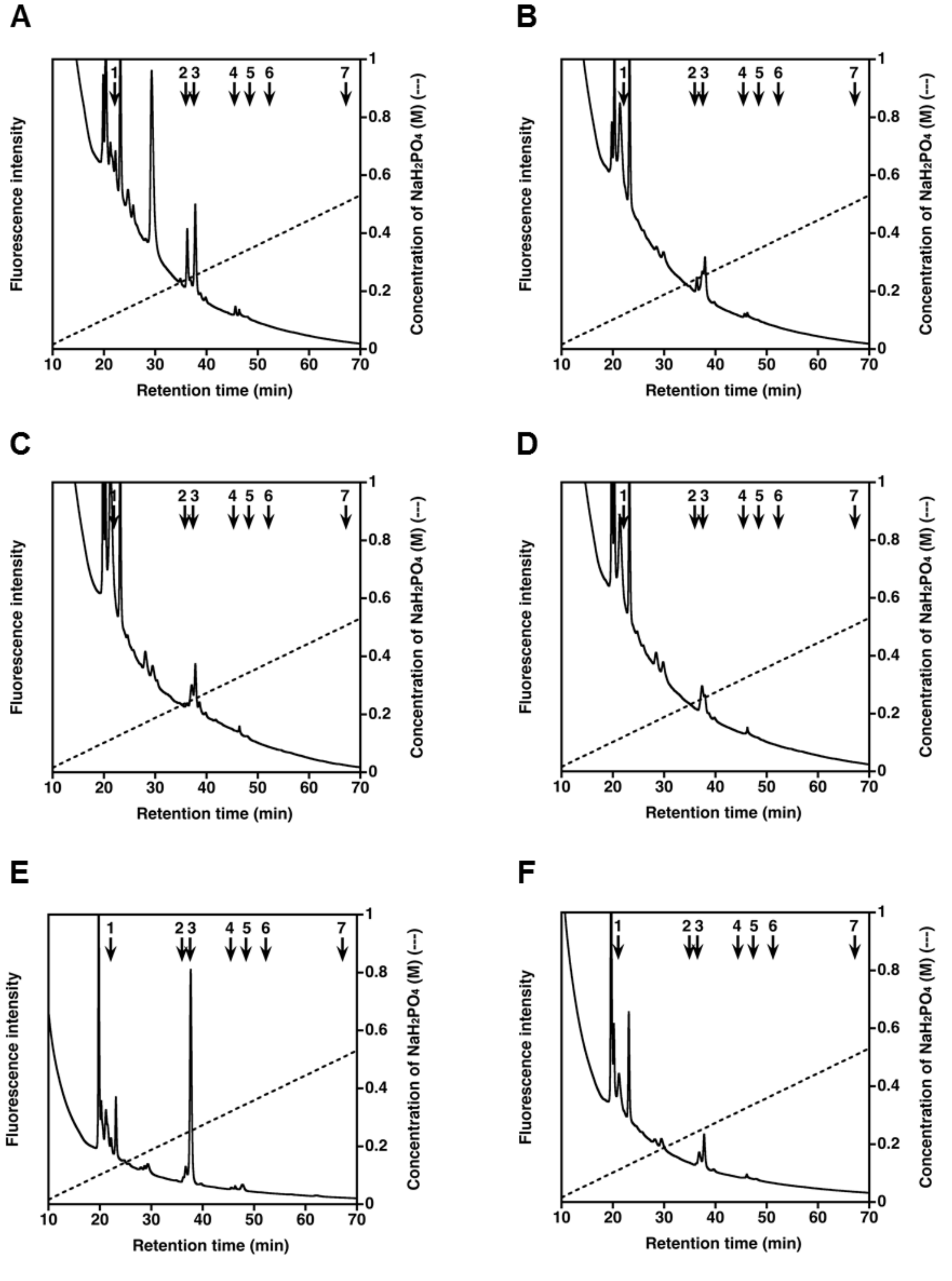


**Fig. S2. HPLC profiles of the GAG-peptide preparations from zebrafish tissues digested with chondroitinase ABC.** The 2AB-derivatives of the yielded CS/DS disaccharides from bone (**A, B**), muscle (**C, D**), and skin (**E, F**) of the wild-type (**A, C, E**) and *b3galt6*$^{-/-}$ zebrafish (**B, D, F**) after digestion with a mixture of chondroitinases ABC and AC-II for analysis of both CS and DS moieties, were separated by anion-exchange HPLC on an amine-bound silica PA-G column using a linear gradient of $NaH_2PO_4$ as indicated by the dashed line. The elution positions of 2AB-labeled CS/DS disaccharide standards are indicated by numbered arrows: 1, ΔHexA-GalNAc; 2, ΔHexA-GalNAc(6S); 3, ΔHexA-GalNAc(4S); 4, ΔHexA(2S)-GalNAc(6S); 5, ΔHexA(2S)-GalNAc(4S); 6, ΔHexA-GalNAc(4S,6S); 7, ΔHexA(2S)-GalNAc(4S,6S). Abbriviations: ΔHexA, GalNAc, 2S, 4S, and 6S represent 4,5-unsaturated hexuronic acid, *N*-acetyl-D-galactosamine, 2-*O*-sulfate, 4-*O*-sulfate, and 6-*O*-sulfate, respectively.

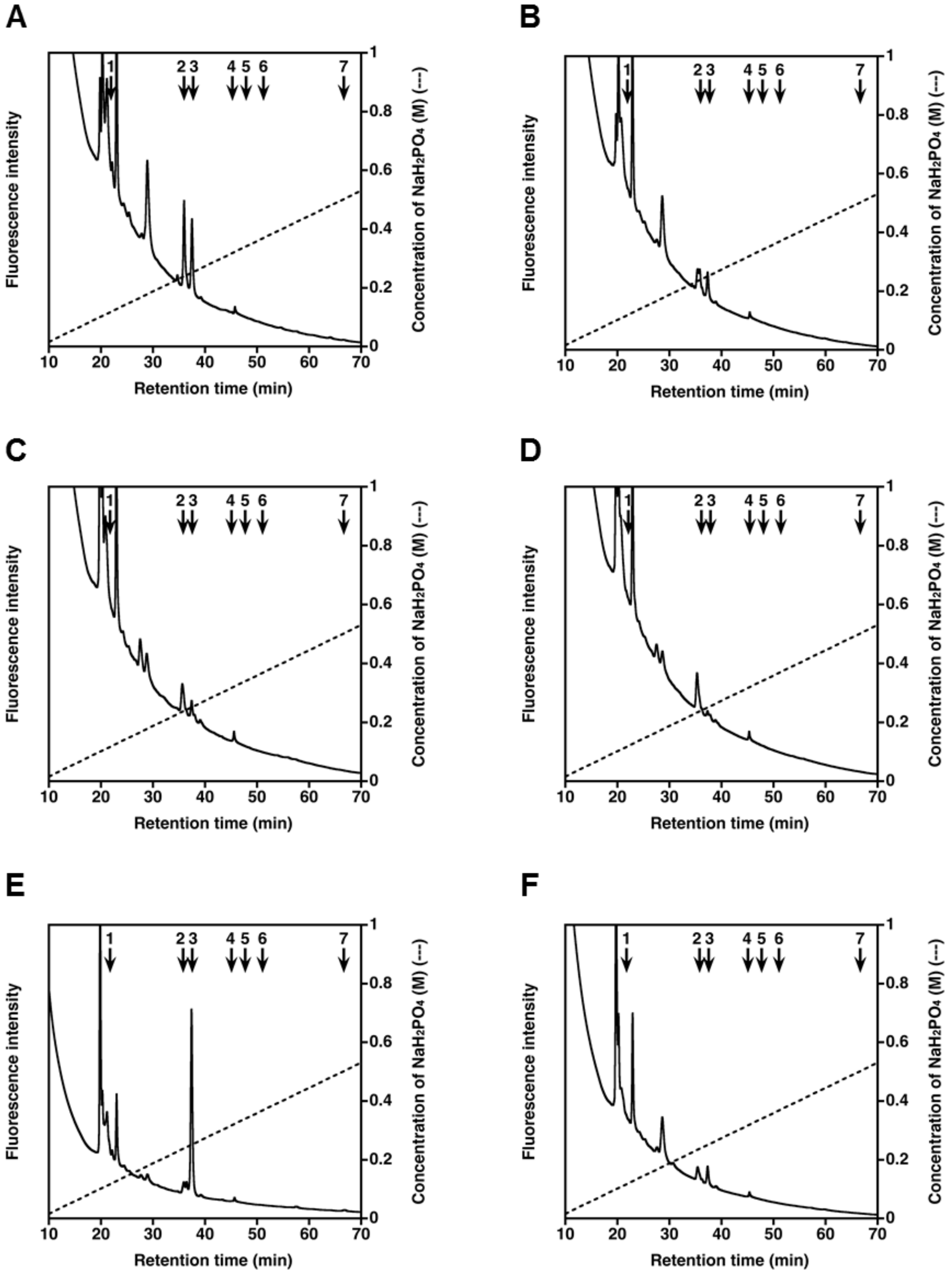

**Fig. S3. HPLC profiles of the GAG-peptide preparations from zebrafish tissues digested with chondroitinase AC.** The 2AB-derivatives of the yielded CS disaccharides from bone (**A, B**), muscle (**C, D**), and skin (**E, F**) of the wild-type (**A, C, E**) and *b3galt6*$^{-/-}$ zebrafish (**B, D, F**) after digestion with a mixture of chondroitinases AC-I and AC-II for analysis of CS moiety, were separated by anion-exchange HPLC on an amine-bound silica PA-G column using a linear gradient of $NaH_2PO_4$ as indicated by the dashed line. The elution positions of 2AB-labeled CS/DS disaccharide standards are indicated by numbered arrows: 1, ΔHexA-GalNAc; 2, ΔHexA-GalNAc(6S); 3, ΔHexA-GalNAc(4S); 4, ΔHexA(2S)-GalNAc(6S); 5, ΔHexA(2S)-GalNAc(4S); 6, ΔHexA-GalNAc(4S,6S); 7, ΔHexA(2S)-GalNAc(4S,6S).

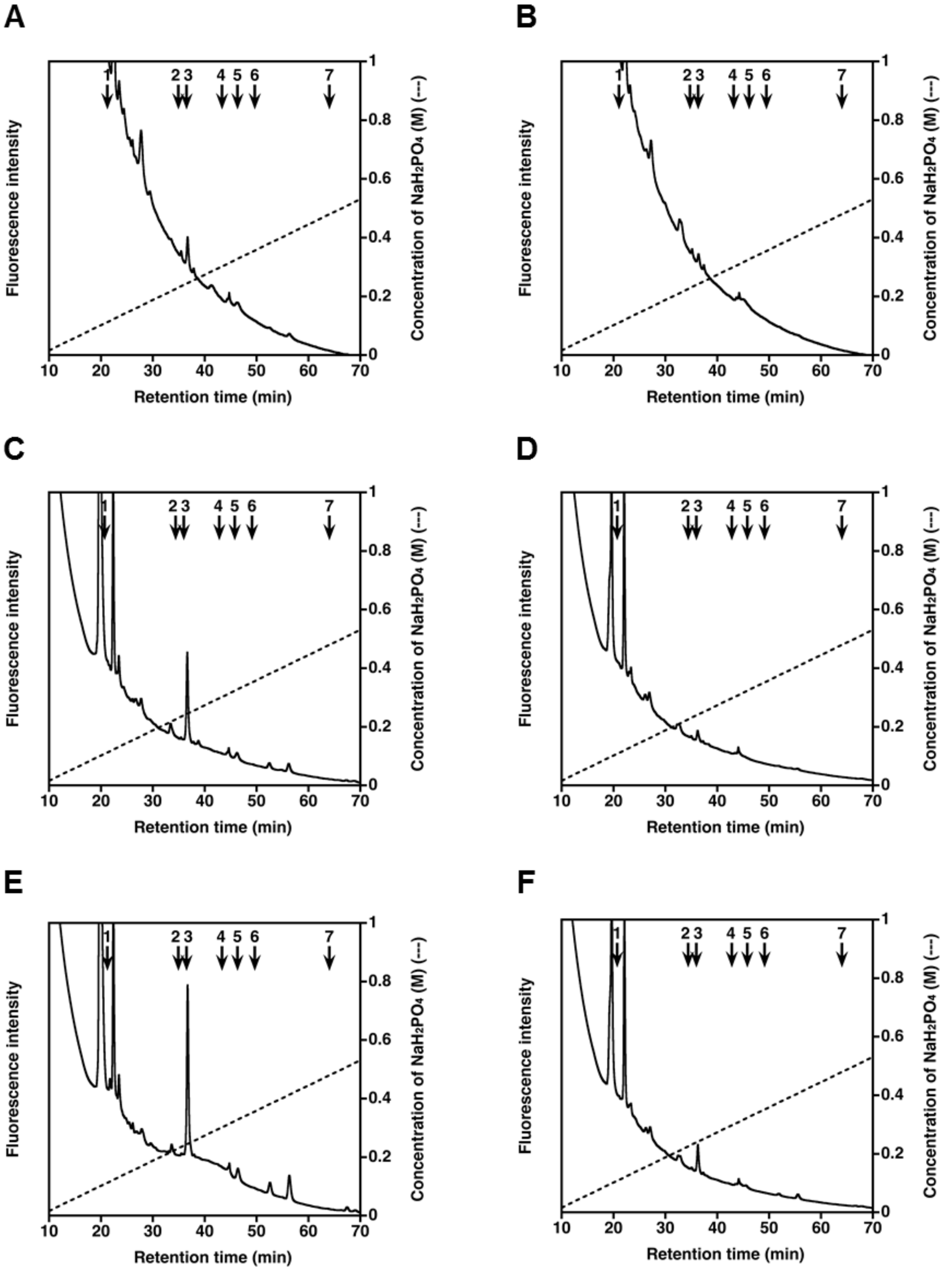


**Fig. S4. HPLC profiles of the GAG-peptide preparations from zebrafish tissues digested with chondroitinase B.** The 2AB-derivatives of the yielded CS disaccharides from bone (**A, B**), muscle (**C, D**), and skin (**E, F**) of the wild-type (**A, C, E**) and *b3galt6*$^{-/-}$ zebrafish (**B, D, F**) after digestion with chondroitinase B for analysis of DS moiety, were separated by anion-exchange HPLC on an amine-bound silica PA-G column using a linear gradient of $NaH_2PO_4$ as indicated by the dashed line. The elution positions of 2AB-labeled CS/DS disaccharide standards are indicated by numbered arrows: 1,

ΔHexA-GalNAc; 2, ΔHexA-GalNAc(6S); 3, ΔHexA-GalNAc(4S); 4, ΔHexA(2S)-GalNAc(6S); 5, ΔHexA(2S)-GalNAc(4S); 6, ΔHexA-GalNAc(4S,6S); 7, ΔHexA(2S)-GalNAc(4S,6S).

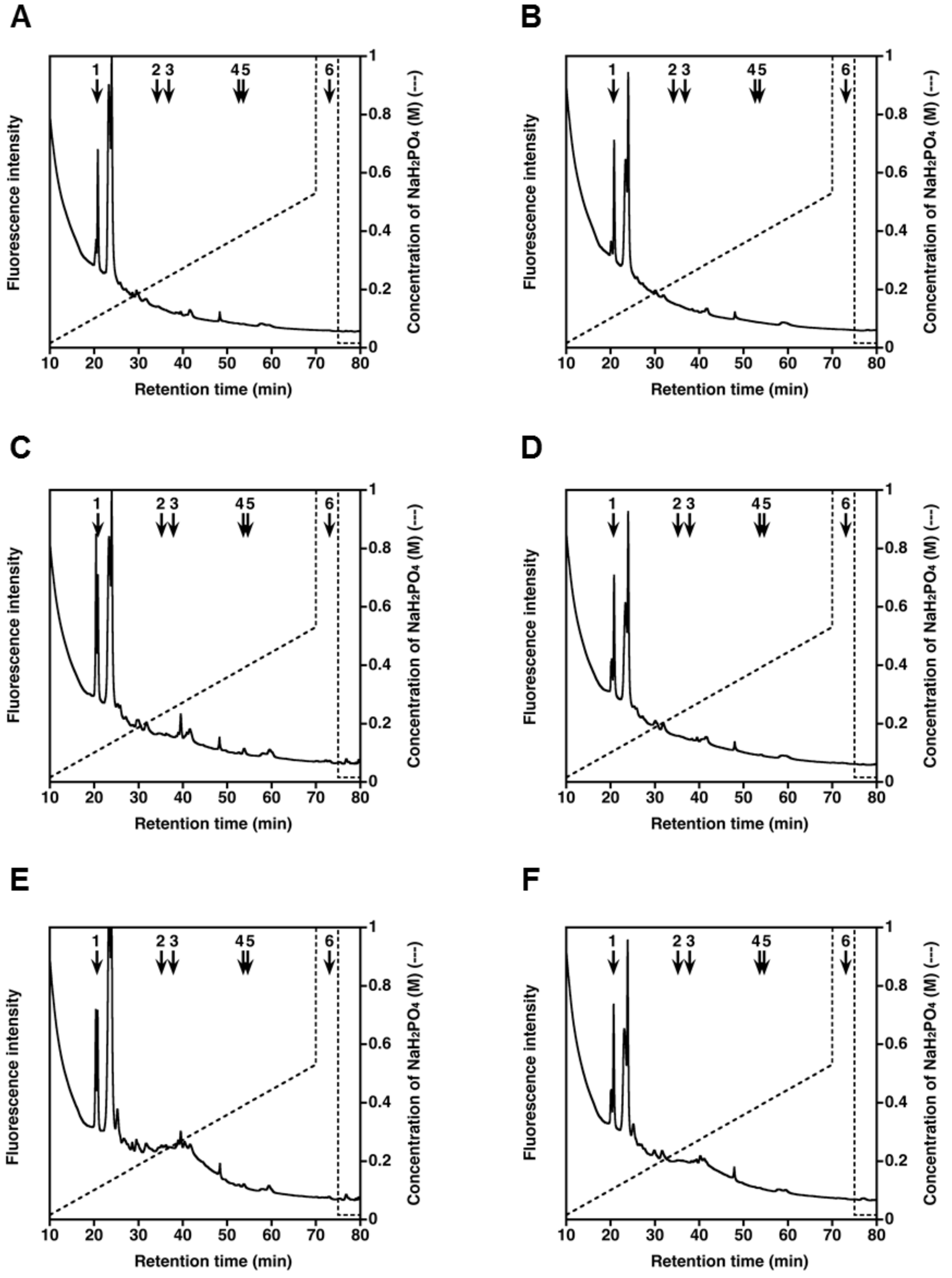


**Fig. S5. HPLC profiles of the GAG-peptide preparations from zebrafish tissues digested with heparinase.** The 2AB-derivatives of the yielded HS disaccharides from bone (**A, B**), muscle (**C, D**), and skin (**E, F**) of the wild-type (**A, C, E**) and *b3galt6*$^{-/-}$ zebrafish (**B, D, F**) after digestion with a mixture of heparinase-I and -III for analysis of HS moiety, were separated by anion-exchange HPLC on an amine-bound silica PA-G column using a linear gradient of $NaH_2PO_4$ as indicated by the dashed line. The elution positions of 2AB-labeled HS disaccharide standards are indicated by numbered arrows: 1, ΔHexA-GlcNAc; 2, ΔHexA-GlcNAc(6S); 3, ΔHexA-GlcN(NS); 4, ΔHexA-GlcN(NS,6S); 5, ΔHexA(2S)-GlcN(NS); 6, ΔHexA(2S)-GlcN(NS,6S). *Abbriviations*: ΔHexA, 4,5-unsaturated hexuronic acid; GlcNAc, *N*-acetyl-D-glucosamine; GlcN, D-glucosamine; 2S, 6S, and NS, 2-*O*-, 6-*O*-, and 2-*N*-sulfate, respectively.

**Table S1. Disaccharide composition of the CS/DS chains.** The GAG fractions prepared from bone, muscle, and skin of the zebrafish were individually digested with a mixture of chondroitinases ABC and AC-II (for the analysis of both CS and DS moieties), and each digest was labeled with 2-AB and analyzed by anion-exchange HPLC (Fig. S2). The amount of resultant disaccharides in each sample was calculated based on the peak area in each chromatogram.

| | **Bone** | | **Muscle** | | **Skin** | |
|---|---|---|---|---|---|---|
| **CS/DS** | Wild-type | *b3galt6*$^{-/-}$ | Wild-type | *b3galt6*$^{-/-}$ | Wild-type | *b3galt6*$^{-/-}$ |
| | *pmol/mg protein* (*mol%*) | | | | | |
| ΔHexA-GalNAc | N.D. | N.D. | N.D. | N.D. | N.D. | N.D. |
| ΔHexA-GalNAc(6S) | 275.8 ± 10.3<br>(35.7 ± 2.0) | 84.7 ± 19.4 [a]<br>(24.0 ± 4.1) | N.D. | N.D. | N.D. | N.D. |
| ΔHexA-GalNAc(4S) | 498.4 ± 26.4<br>(64.3 ± 2.0) | 261.4 ± 0.6 [a]<br>(76.0 ± 4.1) | 9.3 ± 0.4<br>(100 ± 0.0) | 5.7 ± 3.1<br>(100 ± 0.0) | 236.2 ± 26.5<br>(100 ± 0.0) | 55.4 ± 15.2 [c]<br>(100 ± 0.0) |
| ΔHexA(2S)-GalNAc(6S) | N.D. | N.D. | N.D. | N.D. | N.D. | N.D. |
| ΔHexA(2S)-GalNAc(4S) | N.D. | N.D. | N.D. | N.D. | N.D. | N.D. |
| ΔHexA-GalNAc(4S,6S) | N.D. | N.D. | N.D. | N.D. | N.D. | N.D. |
| ΔHexA(2S)-GalNAc(4S,6S) | N.D. | N.D. | N.D. | N.D. | N.D. | N.D. |
| **Total CS/DS disaccharide** | 774.2 ± 19.9 | 346.1 ± 20.0 [c] | 9.3 ± 0.4 | 5.7 ± 3.1 | 236.2 ± 26.5 | 55.4 ± 15.2 [c] |

*Abbreviations*: ΔHexA, 4,5-unsaturated hexuronic acid; GalNAc, *N*-acetyl-D-galactosamine; 2S, 4S, and 6S, 2-*O*-, 4-*O*-, and 6-*O*-sulfate, respectively.

ND, not detected (<1 pmol/mg protein). [a], $p<0.001$ (Student's t-test, vs wild-type); [b], $p<0.0002$; [c], $p<0.005$

**Table S2. Disaccharide composition of the CS moiety of the CS/DS chains.** The GAG fractions prepared from bone, muscle, and skin of the zebrafish were individually digested with a mixture of chondroitinases AC-I and AC-II (for the specific analysis of CS moiety), and each digest was labeled with 2-AB and analyzed by anion-exchange HPLC (Fig. S3). The amount of resultant disaccharides in each sample was calculated based on the peak area in each chromatogram.

| | **Bone** | | **Muscle** | | **Skin** | |
|---|---|---|---|---|---|---|
| **CS** | Wild-type | *b3galt6*$^{-/-}$ | Wild-type | *b3galt6*$^{-/-}$ | Wild-type | *b3galt6*$^{-/-}$ |
| | *pmol/mg protein* (*mol%*) | | | | | |
| ΔHexA-GalNAc | N.D. | N.D. | N.D. | N.D. | N.D. | N.D. |
| ΔHexA-GalNAc(6S) | 403.6 ± 83.6 (55.2 ± 1.6) | 126.8 ± 2.3 [a] (45.3 ± 0.2 [b]) | N.D. | N.D. | 8.9 ± 4.5 (5.4 ± 2.8) | 18.3 ± 10.0 (31.3 ± 15.7) |
| ΔHexA-GalNAc(4S) | 324.8 ± 65.8 (44.8 ± 1.6) | 153.2 ± 2.6 (54.7 ± 0.2 [b]) | 2.1 ± 0.4 (100 ± 0.0) | N.D. | 166.2 ± 25.5 (94.6 ± 2.8) | 29.6 ± 4.8 [c] (68.7 ± 15.7) |
| ΔHexA(2S)-GalNAc(6S) | N.D. | N.D. | N.D. | N.D. | N.D. | N.D. |
| ΔHexA(2S)-GalNAc(4S) | N.D. | N.D. | N.D. | N.D. | N.D. | N.D. |
| ΔHexA-GalNAc(4S,6S) | N.D. | N.D. | N.D. | N.D. | N.D. | N.D. |
| ΔHexA(2S)-GalNAc(4S,6S) | N.D. | N.D. | N.D. | N.D. | N.D. | N.D. |
| **Total CS disaccharide** | 728.4 ± 148.1 | 279.9 ± 4.8 [a] | 2.1 ± 0.4 | —— | 175.1 ± 25.3 | 47.9 ± 13.5 [d] |

*Abbreviations*: ΔHexA, 4,5-unsaturated hexuronic acid; GalNAc, *N*-acetyl-D-galactosamine; 2S, 4S, and 6S, 2-*O*-, 4-*O*-, and 6-*O*-sulfate, respectively.

ND, not detected (<0.1 pmol/mg protein). [a], p<0.05 (Student's t-test, vs wild-type); [b], p<0.005; [c], p<0.01; [d], p<0.02

**Table S3. Disaccharide composition of the DS moiety of the CS/DS chains.** The GAG fractions prepared from bone, muscle, and skin of the zebrafish were individually digested with chondroitinase B (for the specific analysis of DS moiety), and each digest was labeled with 2-AB and analyzed by anion-exchange HPLC (Fig. S4). The amount of resultant disaccharides in each sample was calculated based on the peak area in each chromatogram.

| | **Bone** | | **Muscle** | | **Skin** | |
|---|---|---|---|---|---|---|
| **DS** | Wild-type | *b3galt6*$^{-/-}$ | Wild-type | *b3galt6*$^{-/-}$ | Wild-type | *b3galt6*$^{-/-}$ |
| | *pmol/mg protein* (*mol%*) | | | | | |
| ΔHexA-GalNAc | N.D. | N.D. | N.D. | N.D. | N.D. | N.D. |
| ΔHexA-GalNAc(6S) | N.D. | N.D. | N.D. | N.D. | N.D. | N.D. |
| ΔHexA-GalNAc(4S) | 50.5 ± 8.1 (100 ± 0.0) | N.D. | 14.5 ± 1.1 (100 ± 0.0) | 4.1 ± 1.0 [a] (100 ± 0.0) | 68.0 ± 7.4 (89.5 ± 0.3) | 32.0 ± 5.5 [b] (95.7 ± 4.3) |
| ΔHexA(2S)-GalNAc(6S) | N.D. | N.D. | N.D. | N.D. | N.D. | N.D. |
| ΔHexA(2S)-GalNAc(4S) | N.D. | N.D. | N.D. | N.D. | 7.9 ± 0.6 (10.5 ± 0.3) | 1.8 ± 1.8 [c] (4.3 ± 4.3) |
| ΔHexA-GalNAc(4S,6S) | N.D. | N.D. | N.D. | N.D. | N.D. | N.D. |
| ΔHexA(2S)-GalNAc(4S,6S) | N.D. | N.D. | N.D. | N.D. | N.D. | N.D. |
| **Total DS disaccharide** | 50.5 ± 8.1 | — | 14.5 ± 1.1 | 4.1 ± 1.0 [a] | 75.9 ± 7.9 | 33.8 ± 6.3 [b] |

*Abbreviations*: ΔHexA, 4,5-unsaturated hexuronic acid; GalNAc, *N*-acetyl-D-galactosamine; 2S, 4S, and 6S, 2-*O*-, 4-*O*-, and 6-*O*-sulfate, respectively. ND, not detected (<0.1 pmol/mg protein). [a], $p<0.005$ (Student's t-test, vs wild-type); [b], $p<0.02$; [c], $p<0.05$

**Table S4. Disaccharide composition of the HS chains.** The GAG fractions prepared from bone, muscle, and skin of the zebrafish were individually digested with a mixture of heparinase-I and heparinase-III (for the specific analysis of the HS moiety), and each digest was labeled with 2-AB and analyzed by anion-exchange HPLC (Fig. S5). The amount of resultant disaccharides in each sample was calculated based on the peak area in each chromatogram.

| | **Bone** | | **Muscle** | | **Skin** | |
|---|---|---|---|---|---|---|
| **HS** | Wild-type | *b3galt6*$^{-/-}$ | Wild-type | *b3galt6*$^{-/-}$ | Wild-type | *b3galt6*$^{-/-}$ |
| | *pmol/mg protein* (*mol%*) | | | | | |
| ΔHexA-GlcNAc | 178.6 ± 19.8<br>(100 ± 0.0) | 137.1 ± 39.7<br>(100 ± 0.0) | 25.6 ± 0.9<br>(83.2 ± 0.3) | 13.5 ± 4.0 [a]<br>(100 ± 0.0 [b]) | 47.0 ± 3.7<br>(85.3 ± 0.3) | 21.0 ± 2.8 [d]<br>(100 ± 0.0 [b]) |
| ΔHexA-GlcNAc(6S) | N.D. | N.D. | N.D. | N.D. | N.D. | N.D. |
| ΔHexA-GlcN(NS) | N.D. | N.D. | 5.2 ± 0.3<br>(16.8 ± 0.3) | N.D. | 8.1 ± 0.8<br>(14.7 ± 0.3) | N.D. |
| ΔHexA-GlcN(NS,6S) | N.D. | N.D. | N.D. | N.D. | N.D. | N.D. |
| ΔHexA(2S)-GlcN(NS) | N.D. | N.D. | N.D. | N.D. | N.D. | N.D. |
| ΔHexA(2S)-GlcN(NS,6S) | N.D. | N.D. | N.D. | N.D. | N.D. | N.D. |
| **Total HS disaccharide** | 178.6 ± 19.8 | 137.1 ± 39.7 | 30.8 ± 1.2 | 13.5 ± 4.0 [c] | 55.1 ± 4.5 | 21.0 ± 2.8 [d] |

*Abbreviations*: ΔHexA, 4,5-unsaturated hexuronic acid; GlcNAc, *N*-acetyl-D-glucosamine; GlcN, D-glucosamine; 2S, 6S, and NS, 2-*O*-, 6-*O*-, and 2-*N*-sulfate, respectively. ND, not detected (<0.1 pmol/mg protein). [a], $p<0.05$ (Student's t-test, vs wild-type); [b], $p<0.0001$; [c], $p<0.02$; [d], $p<0.005$.

**Table S5. The level of digested GAG disaccharides from wild-type and *b3galt6*$^{-/-}$ (cmg22) zebrafish tissues.**

| | *pmol/mg protein (Average ± SE)* | | | |
|---|---|---|---|---|
| **Bone** | CS/DS | CS | DS | HS |
| Wild-type | 884.1 ± 327.2 | 782.7 ± 293.2 | 30.7 ± 21.2 | 233.4 ± 159.3 |
| *b3galt6*$^{-/-}$ | 307.0 ± 136.6 | 336.6 ± 135.5 | 17.9 ± 11.9 | 43.5 ± 19.3 |
| **Muscle** | | | | |
| Wild-type | 48.0 ± 5.5 | 20.4 ± 4.4 | 17.6 ± 2.6 | 113.8 ± 76.0 |
| *b3galt6*$^{-/-}$ | 20.8 ± 4.9* | 14.2 ± 4.1 | 7.0 ± 1.7* | 8.3 ± 2.2 |
| **Skin** | | | | |
| Wild-type | 1,066 ± 80.8 | 914.8 ± 76.0 | 74.1 ± 16.6 | 56.0 ± 15.2 |
| *b3galt6*$^{-/-}$ | 152.9 ± 26.3*** | 123.42 ± 20.6*** | 18.1 ± 4.6* | 17.1 ± 5.4* |

Student's t-test (vs wild-type) (n=5), *p<0.05, **p<0.01, ***p<0.0001. SE: standard error.

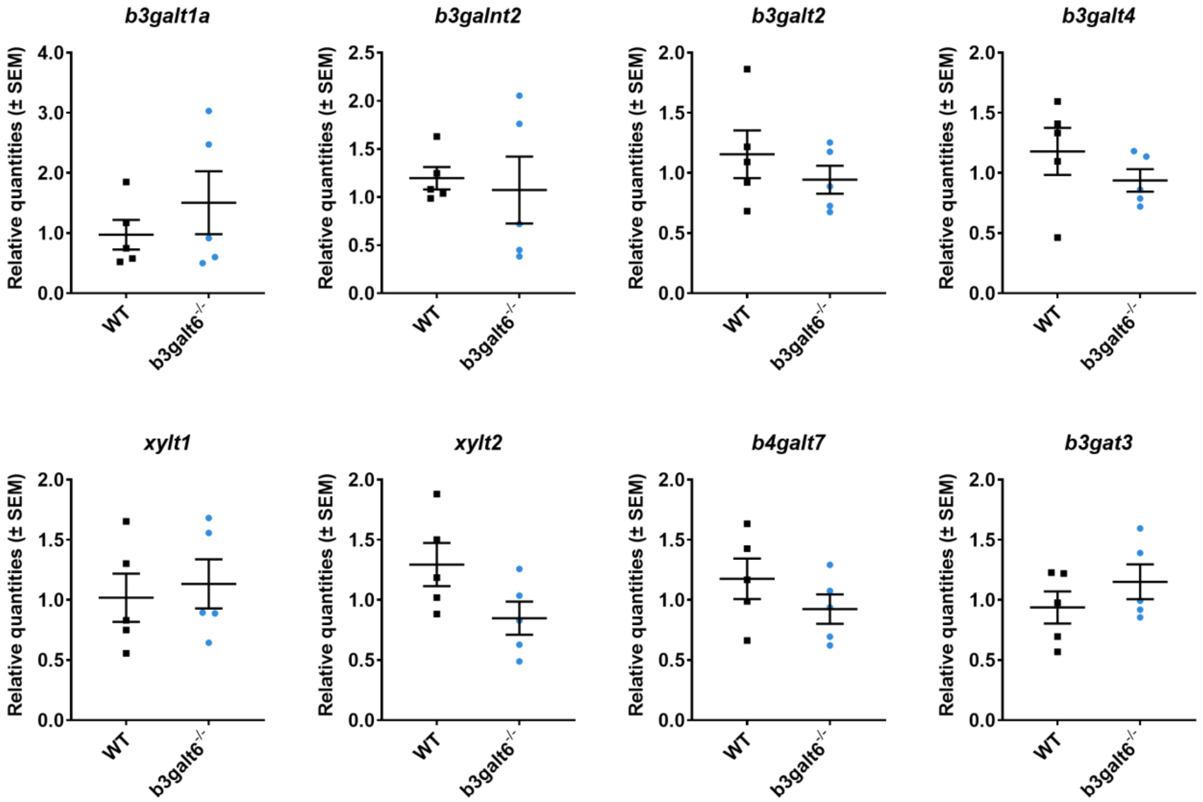


**Fig. S6 mRNA expression levels of *b3galt* family members and proteoglycan linkage genes.** RT-qPCR analysis of the *b3galt* family members (*b3galt1a, b3galnt2, b3galt2* and *b3galt4*) and other proteoglycan linkage genes (*xylt1, xylt2, b4galt7* and *b3gat3*) excluding *b3galt6*. Their mRNA expression levels were not significantly different in adult *b3galt6*$^{-/-}$ (cmg20) mutants compared to the expression levels in adult WT siblings. Data are expressed as mean ± SEM (n=5). An unpaired t-test was used to determine significance. Respective P-values: 0.3840 (*b3galt1a*), 0.7470 (*b3galnt2*), 0.3844 (*b3galt2*), 0.2984 (*b3galt4*), 0.6994 (*xylt1*), 0.0840 (*xylt2*), 0.9240 (*b4galt7*) and 0.3120 (*b3gat3*).

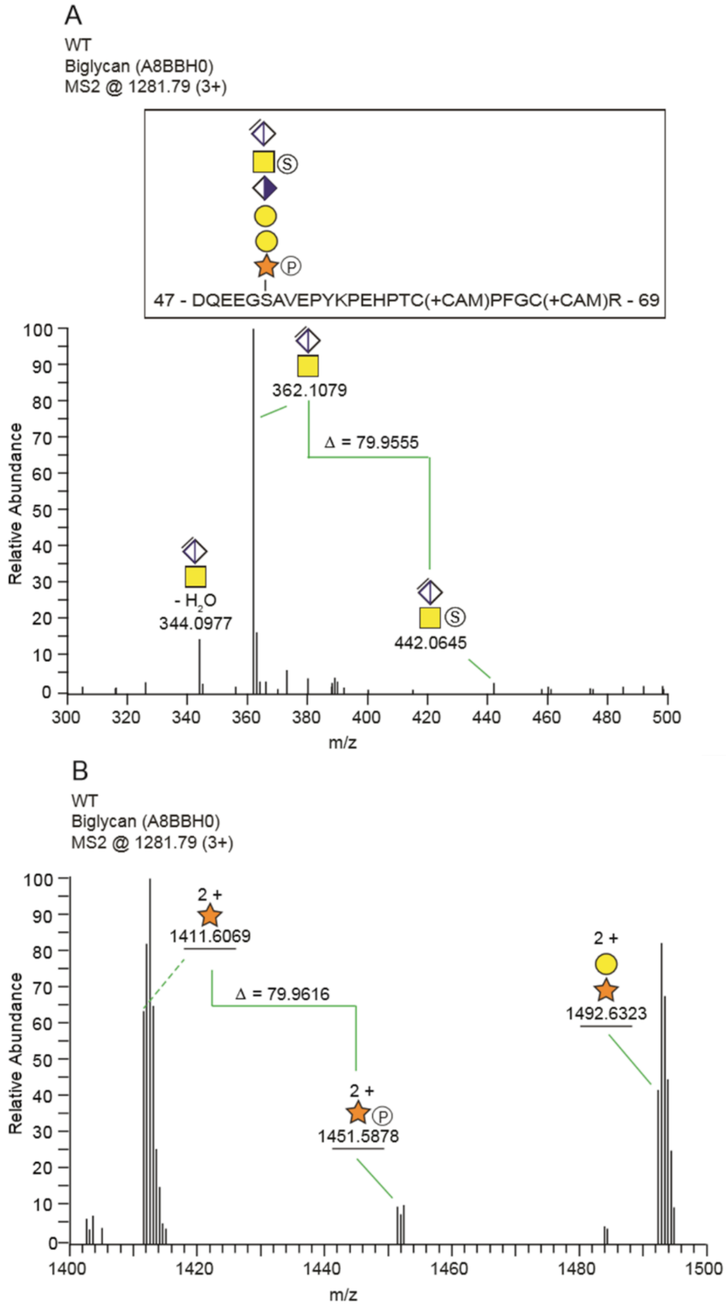


**Fig. S7. Structural analysis of the biglycan CS-linkage region in WT zebrafish.** (**A**) The MS2 fragment spectrum displayed a mass shift of 79.9555 Da between *m/z* = 362.1079; 1+ and 442.0645; 1+, demonstrating the presence of a sulfate group on the GalNAc residue. (**B**) In the higher mass-range (*m/z* 1400 - 1500), the spectrum displayed a mass shift of 79.9616 Da between *m/z* 1411.6069; 2+ and 1492.6323; 2+, demonstrating the presence of a phosphate group.

**Table S6. List of all HS- and CS glycopeptides identified in WT and *b3galt6*$^{-/-}$ zebrafish.**

| Protein name[a] | Uniprot /NCBI | Peptide sequence | Modification[b] | m / z; z | Accuracy (ppm)[c] |
|---|---|---|---|---|---|
| **WT** | | | | | |
| Aggrecan core protein, isoform X1 | XP_021326217.1 | $^{987}$GTSGYPNHPSGEVSENIQR$^{1005}$ | CS, hexasaccharide | 1008.08; 3+ | + 1.72 |
| Aggrecan core protein, isoform X3 | XP_686182.6 | $^{1076}$LGSGFTDLTEQSR$^{1088}$ | CS, hexasaccharide | 1202.49; 3+ | + 2.37 |
| Agrin | A0A0R4IJ40 | $^{561}$PGVCDEDCGSGGSGSGSETCEQDR$^{584}$ | HS, tetrasaccharide | 1039.69; 3+ | - 0.34 |
| Biglycan | A8BBH0 | $^{47}$DQEEGSAVEPYKPEHPTCPFGCR$^{69}$ | CS, hexasaccharide + S and P | 1281.79; 3+ | + 2.33 |
| Collagen type XVIII, alpha 1 | B0S8G4 | $^{764}$DMEGSGFDMDSVR$^{776}$ | HS, tetrasaccharide | 1030.37; 2+ | - 0.037 |
| Collagen type XV-B alpha 1 chain | A0A0A1I9R6 | $^{243}$SGDGDYDDEEEHAR$^{257}$ | HS, tetrasaccharide + S / P | 1144.86; 2+ | + 0.47 |
| Ephiphycan | B0S5X0 | $^{65}$RVDPFEGIESSHSASLEEK$^{83}$ | CS, hexasaccharide | 1064.08; 3+ | + 0.070 |
| Osteopontin | A1KXD6 | $^{166}$AVLQVHDLEDGNTSTSEIDNIER$^{185}$ | CS, hexasaccharide + S / P | 1281.84; 3+ | + 1.32 |
| Osteopontin | A1KXD6 | $^{190}$QAALGAQEIVPVGDQAATEEGASAR$^{211}$ | CS, hexasaccharide + S / P | 1171,49;3+ | + 0.26 |
| Sushi-repeat containing protein | Q58ED3 | $^{21}$YDGSGYWPYTDDEDVT$^{35}$ | HS, tetrasaccharide | 1276.48; 2+ | + 1.51 |
| Syndecan-3 | D1DFN3 | $^{99}$ESGLGQDVDSK$^{109}$ | CS, hexasaccharide + S / P | 1104.38; 2+ | + 0.044 |
| [a]Uncharacterized protein si:ch73-306e8.2 isoform X1 | XP_00133717.3 | $^{98}$SESEHSGSGDVSSEK$^{113}$ | CS, hexasaccharide | 1341.46; 2+ | + 2.48 |
| ***b3galt6***$^{-/-}$ | | | | | |
| Aggrecan core protein, isoform X2 | XP_021329528.1 | $^{851}$AFGEQELSGFY$^{861}$ | CS, pentasaccharide | 1039.89; 3+ | + 1.43 |
| Aggrecan core protein, isoform X3 | XP_686182.6 | $^{1076}$LGSGFTDLTEQSR$^{1088}$ | CS, pentasaccharide | 1121.46; 2+ | + 2.01 |
| Biglycan | A8BBH0 | $^{47}$DQEEGSAVEPYKPEHPTCPFGCR$^{69}$ | CS, pentasaccharide + 2 x S / P | 1227.77; 3+ | + 1.54 |

[a] Indicate novel PG identified in the present study.

[b] Mascot searches identified glycan structures either without, or with, sulfate (S)- and/or phosphate (P) modifications. The distinction of potential sulfate (79.9663 Da) and phosphate (79.9568 Da) modifications are not provided by the Mascot searches. Structures with modifications are therefore marked S / P to denote the

ambiguity of the modification. In the case of WT biglycan, the distinction between S and P was made by manually examining the MS2 spectra in close detail (Fig. S7).

[c] The accuracy denotes the difference between the theoretical and the measured mass for each CS/HS - glycopeptide. For glycopeptides which contain modifications (i.e. sulfate- or phosphate) the theoretical mass is calculated based on the assumption of sulfate modification(s).

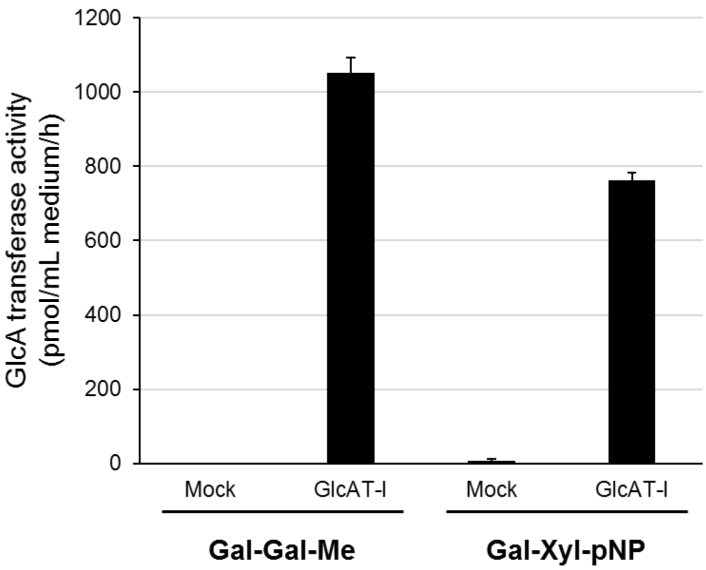


**Fig. S8. GlcA transferase activity of recombinant B3GAT3/GlcAT-I toward Gal-Gal and Gal-Xyl *in vitro*.** The experiment was performed in triplicate. Gal, galactose; Xyl, Xylose; Me, methyl; pNP, *p*-nitrophenyl.

**Table S7. Developmental table comparing *b3galt6* WT and mutant siblings.**

| Structure | *b3galt6*$^{+/+}$ | | | | | *b3galt6*$^{-/-}$ | | | | |
|---|---|---|---|---|---|---|---|---|---|---|
| **Name (abbreviation)** | **1** | **2** | **3** | **4** | **5** | **1** | **2** | **3** | **4** | **5** |
| *Head* | | | | | | | | | | |
| Ethmoid plate (ep) | + | + | + | + | + | + | + | + | + | + |
| Lamina orbitonasalis (lon) | + | + | + | + | + | ± | ± | ± | ± | ± |
| Taenia marginalis anterior (tma) | + | + | + | + | + | - | ± | - | + | + |
| Epiphysial bar (epb) | + | + | + | + | + | ± | + | ± | + | ± |
| Sclerotic cartilage | + | + | + | + | + | ± | ± | ± | + | ± |
| Taenia marginalis posterior (tmp) | + | + | + | + | + | + | + | + | + | + |
| Anterior basicapsular commissure (abc) | + | + | + | + | + | + | + | + | + | + |
| Basicapsular fenestra (b) | + | + | + | + | + | + | + | + | + | + |
| Posterior basicapsular commissure (pbc) | + | + | + | + | + | + | + | + | + | + |
| Auditory capsule (ac) | + | + | + | + | + | ± | ± | ± | + | ± |
| Tectum synoticum (ts) | + | + | + | ± | + | ± | ± | ± | ± | ± |
| Basioccipital (boc) | + | + | + | + | + | + | + | + | + | + |
| *Pectoral fin* | | | | | | | | | | |
| Scapula (s) | ± | ± | ± | ± | ± | - | - | - | - | - |
| Scapulacoracoid (sc) | + | + | + | + | + | + | + | + | + | + |
| Precoracoid process (prp) | + | + | + | + | + | + | + | + | + | + |
| Postcoracoid process (prp) | + | + | + | + | + | + | + | + | + | + |
| Fin disk (fd) | + | + | + | + | + | - | ± | - | ± | - |
| Central separation zone 1 (csz 1) | + | + | + | + | + | - | - | - | - | - |
| *Caudal fin* | | | | | | | | | | |
| Neural arch/spine on PU3 (na/s pu3) | - | - | ± | - | ± | - | - | - | - | - |
| Neural arch/spine on PU2 (na/s pu3) | ± | + | ± | ± | ± | - | - | - | ± | - |
| Vestigial neural arch/spine on PU1+U (vna/s) | + | + | - | - | - | - | - | - | - | - |
| Epural (e) | + | + | + | + | + | - | - | - | ± | ± |
| Opistural cartilage (opc) | + | + | + | + | + | - | - | - | ± | ± |
| Haemal arch/spine on PU3 (ha/s pu3) | ± | +* | ± | ± | ± | - | - | - | ± | ± |
| Haemal arch/spine on PU2 (ha/s pu2) | + | + | + | + | + | ± | ± | ± | + | ±* |
| Parhypural (phy) | + | + | + | + | + | ±* | + | ± | + | +[2] |
| Hypural 1 (hy1) | + | + | + | + | + | + | + | + | +* | + |
| Hypural 2 (hy2) | + | + | + | + | + | + | + | + | + | + |
| Hypural 3$^{+}$ (hy3$^{+}$) | + | + | +[1] | + | + | + | + | + | + | + |
| Hypural 4 (hy4) | +[1] | + | + | + | + | + | + | + | + | + |
| Hypural 5 (hy5) | + | + | + | + | + | - | - | - | ± | - |

+ present (completely developed), ± developing, - absent

[1] extra cartilaginous element present

[2] base developed separated from hypural 1

* malformed

**Table S8. Mineralization status of individual bones compared between *b3galt6* WT and mutant siblings.**

| Structure | *b3galt6*$^{+/+}$ | | | | | | | | | | *b3galt6*$^{-/-}$ | | | | | | | | | |
|---|---|---|---|---|---|---|---|---|---|---|---|---|---|---|---|---|---|---|---|---|
| **Name** | **1** | **2** | **3** | **4** | **5** | **6** | **7** | **8** | **9** | **10** | **1** | **2** | **3** | **4** | **5** | **6** | **7** | **8** | **9** | **10** |
| *Head-olfactory region* | | | | | | | | | | | | | | | | | | | | |
| Lateral ethmoid | ± | + | + | + | + | + | + | + | + | + | + | ± | ± | + | + | ± | + | + | + | + |
| Vomer | ± | ± | ± | ± | ± | ± | ± | ± | ± | + | ± | - | ± | - | - | ± | ± | ± | ± | ± |
| Supraethmoid | ± | - | ± | ± | ± | ± | ± | ± | ± | + | ± | - | ± | - | ± | - | ± | ± | - | ± |
| Kinethmoid | ± | ± | ± | ± | ± | ± | ± | - | ± | + | - | - | - | - | ± | - | - | ± | - | ± |
| Ethmoid | - | - | - | - | - | - | - | - | - | ± | - | - | - | - | - | - | - | - | - | - |
| *Head-orbital region* | | | | | | | | | | | | | | | | | | | | |
| Parasphenoid | + | + | + | + | + | + | + | + | + | + | + | +* | + | +* | + | +* | +* | +* | + | +* |
| Pterosphenoid | + | + | + | + | + | ± | ± | + | + | + | + | ± | ± | + | + | ± | + | + | ± | + |
| Infraorbital 1 | ± | ± | ± | ± | ± | ± | ± | ± | ± | + | ± | - | ± | - | ± | - | - | ± | - | ± |
| Orbitosphenoid | ± | + | ± | ± | + | ± | ± | + | ± | + | + | ± | - | ± | ± | - | + | + | ± | + |
| Frontal | ± | ± | + | + | + | ± | ± | + | + | + | ± | ± | ± | ± | + | - | ± | ± | ± | ± |
| Nasal | - | - | - | ± | - | - | - | - | - | - | - | - | - | - | - | - | - | - | - | - |
| Supraorbital | - | - | ± | ± | - | - | - | + | - | + | ± | - | - | - | + | - | ± | ± | - | ± |
| *Head-otic region* | | | | | | | | | | | | | | | | | | | | |
| Parietal | - | - | - | - | - | - | - | - | - | ± | - | - | - | - | - | - | - | - | - | - |
| Prootic | + | + | + | + | + | + | + | + | + | + | + | + | ± | + | + | ± | + | + | ± | + |
| Sphenotic | + | + | + | + | ± | + | ± | + | ± | + | ± | ± | ± | ± | + | ± | + | + | ± | + |
| Posttemporal | ± | ± | ± | + | ± | + | ± | ± | ± | + | ± | - | - | ± | ± | - | ± | ± | - | - |
| Pterotic | + | + | + | + | + | + | + | + | + | + | ± | ± | ± | ± | ± | - | ± | ± | ± | ± |
| *Head-occipital region* | | | | | | | | | | | | | | | | | | | | |
| Exoccipital | + | + | + | + | + | + | + | + | + | + | + | + | + | + | + | ± | + | + | + | + |
| Basioccipital | + | + | + | + | + | + | + | + | + | + | + | ± | + | + | + | ± | + | + | + | + |
| Supraoccipital | + | + | ± | + | ± | ± | ± | + | ± | + | ± | ± | ± | ± | ± | - | - | ± | ± | ± |
| *Head-mandibular arch* | | | | | | | | | | | | | | | | | | | | |
| Maxilla | + | + | + | + | + | + | + | + | + | + | + | + | + | + | + | ± | + | + | + | + |
| Premaxilla | + | + | + | + | + | + | + | + | + | + | + | + | + | + | + | ± | + | + | + | + |
| Dentary | + | + | + | + | + | + | + | + | + | + | + | + | + | + | + | ± | + | + | + | + |
| Retroarticular | + | + | + | + | + | + | + | + | + | + | + | + | + | + | + | ± | + | + | + | + |

| | | | | | | | | | | | | | | | | | | | | |
|---|---|---|---|---|---|---|---|---|---|---|---|---|---|---|---|---|---|---|---|---|
| Anguloarticular | + | + | + | + | + | + | + | + | + | + | + | + | + | + | + | + | + | + | + | + |
| Coronomeckelian | + | + | + | + | + | + | + | + | + | + | + | ± | ± | ± | ± | + | ± | + | ± | + |
| *Head-palatoquadrate arch* | | | | | | | | | | | | | | | | | | | | |
| Entopterygoid | + | + | + | + | + | + | + | + | + | + | + | + | + | + | + | - | + | + | ± | + |
| Quadrate | + | + | + | + | + | + | + | + | + | + | + | + | + | ± | + | ± | + | + | + | + |
| Ectopterygoid | + | + | + | ± | + | + | + | + | + | + | + | ± | + | + | + | ± | + | + | + | + |
| Metapterygoid | + | + | + | + | + | + | + | + | + | + | + | + | + | + | + | ± | + | + | + | + |
| Palatine | + | + | + | + | + | + | + | + | + | + | ± | - | ± | ± | ± | - | ± | ± | ± | ± |
| *Head-Hyoid arch* | | | | | | | | | | | | | | | | | | | | |
| Branchiostegal rays (1-3) | + | + | + | + | + | + | + | + | + | + | +* | +* | + | +* | + | +* | +* | + | +* | +* |
| Hyomandibula | + | + | + | + | + | + | + | + | + | + | + | + | + | + | + | + | + | + | + | + |
| Symplectic | + | + | + | + | + | + | + | + | + | + | + | + | + | + | + | + | + | + | + | + |
| Urohyal | + | + | + | + | + | + | + | + | + | + | + | + | + | + | + | + | + | + | + | + |
| Ceratohyal | + | + | + | + | + | + | + | + | + | + | + | + | + | + | + | ± | + | + | + | + |
| Hypohyal (ventral) | + | + | + | + | + | + | + | + | + | + | + | +* | + | + | + | ± | + | + | + | + |
| Hypohyal (dorsal) | ± | ± | - | ± | + | ± | - | ± | ± | + | - | - | ± | ± | + | - | - | - | ± | ± |
| Epihyal | + | + | + | + | + | + | + | + | + | + | + | ± | + | - | - | ± | + | + | + | + |
| basihyal | + | + | + | + | + | + | + | + | + | + | + | + | + | + | + | - | ± | + | ± | + |
| *Head-branchial arches* | | | | | | | | | | | | | | | | | | | | |
| Ceratobranchial 1 | + | + | + | + | + | + | + | + | + | + | + | + | + | ± | + | - | + | + | + | + |
| Ceratobranchial 2 | + | + | + | + | + | + | + | + | + | + | + | + | + | ± | + | ± | ± | + | ± | ± |
| Ceratobranchial 3 | + | + | + | + | + | + | + | + | + | + | + | ± | ± | ± | ± | ± | ± | ± | ± | ± |
| Ceratobranchial 4 | + | + | + | + | + | + | + | + | + | + | + | + | + | ± | + | ± | + | + | ± | ± |
| Ceratobranchial 5 | + | + | + | + | + | + | + | + | + | + | + | + | + | + | + | + | + | + | + | + |
| Epibranchial 1 | + | + | + | + | + | + | + | + | + | + | ± | ± | + | + | ± | ± | ± | + | ± | + |
| Epibranchial 2 | + | + | + | + | + | + | + | ± | ± | + | ± | ± | ± | ± | ± | - | ± | + | ± | + |
| Epibranchial 3 | + | + | + | + | + | + | + | ± | + | + | ± | ± | ± | ± | ± | ± | ± | ± | ± | + |
| Epibranchial 4 | + | + | + | + | + | + | + | + | + | + | + | + | + | + | ± | ± | ± | + | ± | + |
| Basibranchial 1 | - | - | - | - | - | - | - | - | - | ± | - | - | - | - | - | - | - | - | - | - |
| Basibranchial 2 | + | ± | + | + | ± | ± | ± | ± | ± | + | ± | ± | + | ± | ± | - | ± | ± | - | ± |
| Basibranchial 3 | - | ± | ± | ± | - | - | - | - | - | + | - | - | - | - | - | - | - | - | - | - |
| Hypobranchial | - | - | - | - | - | - | - | - | - | - | - | - | - | - | - | - | - | - | - | - |
| *Head-Opercula* | | | | | | | | | | | | | | | | | | | | |

| | | | | | | | | | | | | | | | | | | | | |
|---|---|---|---|---|---|---|---|---|---|---|---|---|---|---|---|---|---|---|---|---|
| Opercle | + | + | + | + | + | + | + | + | + | + | + | + | + | + | + | + | + | + | + | + |
| Interopercle | + | + | + | + | + | + | + | + | + | + | + | + | + | + | + | ± | + | + | + | + |
| Subopercle | + | + | + | + | + | + | + | + | + | + | + | + | + | ± | + | ± | +* | + | + | +* |
| Preopercle | + | + | + | + | + | + | + | + | + | + | + | + | + | + | + | - | ± | ± | ± | ± |
| *Pectoral fin girdle* | | | | | | | | | | | | | | | | | | | | |
| Supracleithrum | + | +* | + | +* | +* | +* | + | + | + | + | +* | + | + | +* | + | ±* | + | + | +* | + |
| Cleithrum | + | + | + | + | + | + | + | + | + | + | + | + | + | + | + | + | + | + | + | + |
| Precoracoid process | - | - | - | ± | - | - | - | - | - | + | - | - | - | - | ± | - | - | - | - | ± |
| scapula | - | - | - | - | - | - | - | - | - | ± | - | - | - | - | ± | - | - | - | - | ± |
| *Caudal fin* | | | | | | | | | | | | | | | | | | | | |
| Preural 3 | + | + | + | + | + | + | + | + | + | + | + | ± | + | + | + | + | +* | - | + | + |
| Neural arch PU3 | + | + | +* | + | +* | + | + | + | +* | + | +* | ±* | +[1] | +* | +[1] | ±*[1] | +* | - | +* | +* |
| Haemal arch PU3 | + | + | + | +[1] | + | +* | +* | +* | +* | +* | +* | ± | +[1] | +*[1] | +*[1] | ± | +*[1] | - | +* | - |
| Preural2 | + | + | + | + | + | + | + | + | + | + | + | + | + | + | + | ± | + | + | + | + |
| Neural arch PU2 | ± | + | + | + | + | +* | + | +* | +* | + | +* | ±[1] | + | ± | +*[1] | ±* | ± | +[1] | ±* | ± |
| Haemal arch PU2 | +* | + | + | + | + | + | + | + | + | + | + | + | + | +* | + | ± | + | +[1] | ± | +[1] |
| Preural1[+] | + | + | + | + | + | + | + | + | + | + | +* | + | + | + | + | ± | + | + | ± | + |
| Vestigial neural arches | + | + | + | + | + | + | + | + | + | + | + | + | + | + | + | + | + | + | + | + |
| Ural 1 | + | + | + | + | + | + | + | + | + | + | + | + | + | + | + | ± | + | + | ± | + |
| Ural 2 | + | + | ± | + | ± | - | + | + | + | + | ± | - | - | ± | +* | - | - | ±* | - | - |
| Parhypural | + | + | + | + | + | + | + | + | + | + | + | + | + | + | + | + | + | + | + | + |
| Epural | ± | ± | ± | ± | ± | ± | ± | ± | ± | + | ± | - | - | - | ±* | - | ±* | ±* | ±* | - |
| Uroneural | + | + | + | +* | + | + | + | + | + | + | + | + | + | +* | +* | + | +* | +[1] | +* | +* |
| Hypural 1 | +* | + | + | + | + | + | + | +* | + | + | +* | + | + | + | + | + | + | + | + | + |
| Hypural 2 | + | + | + | + | + | + | + | +* | + | + | + | + | + | + | + | + | + | + | + | + |
| Hypural 3 | + | + | + | + | + | + | + | +* | + | + | + | + | + | + | + | + | + | +[1] | + | + |
| Hypural 4 | + | + | + | + | + | + | + | + | + | + | + | + | + | + | + | + | + | + | + | + |
| Hypural 5 | + | +[1] | + | + | + | + | + | + | + | + | + | + | + | + | + | + | + | + | + | + |

151 + present (mineralization well underway), ± started to mineralize, - not yet mineralized

152 [1] extra bony element present

153 * malformed

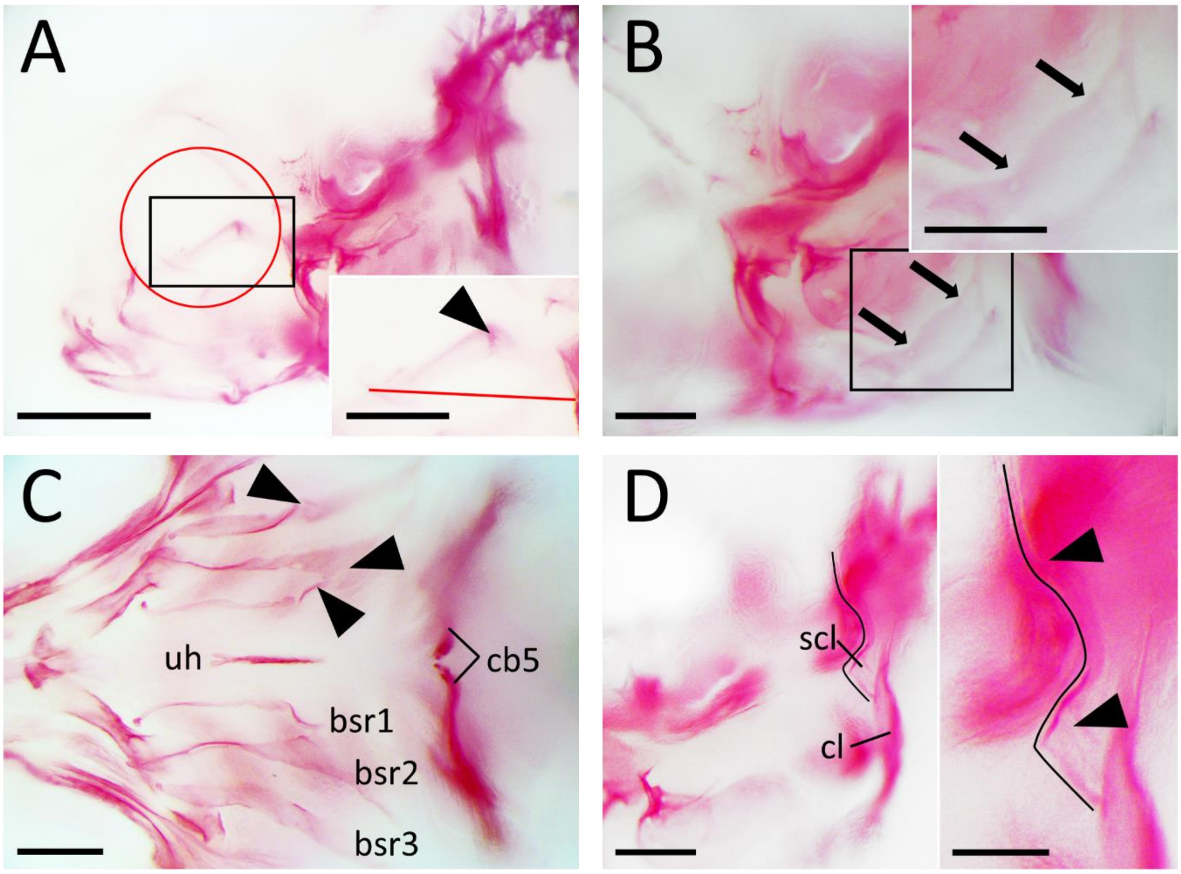


**Fig. S9. Malformations of bones in the head of 20 dpf *b3galt6*$^{-/-}$ zebrafish.** All images are oriented anterior to the left, posterior to the right, dorsal to the top and ventral to the bottom. (**A**) Mutant zebrafish stained with Alizarin red for mineralized bones could have a malformed parasphenoid bone. The malformation was present as a kink in the bone (arrowhead in the magnification) to the dorsal side. The red line indicates where a normally formed parasphenoid would be present. The kink in the bone is always located in the orbital region behind the eye. The position of the eye is indicated by the red circle. Scale bars: 500 µm and 250 µm in the magnification. (**B**) The subopercle could be malformed showing a waved dorsal bone edge (arrows) in mutant zebrafish. Scale bars: 200 µm in both images. (**C**) The branchiostegal rays (bsr1-3) showed kinks, enlarged areas of intramembranous bone and intramembranous bony outgrowths (arrowheads) in mutant zebrafish. Further abbreviations: cb5, ceratobranchial 5, uh, urohyal. Scale bar: 200 µm. (**D**) Both WT and mutant sibling zebrafish could have a malformed supracleithrum (scl), with a kink both in the dorsal and ventral part of the bone (arrowhead in magnification). Further abbreviations: cl, cleithrum. Scale bars: 200 µm and 100 µm in the magnification.

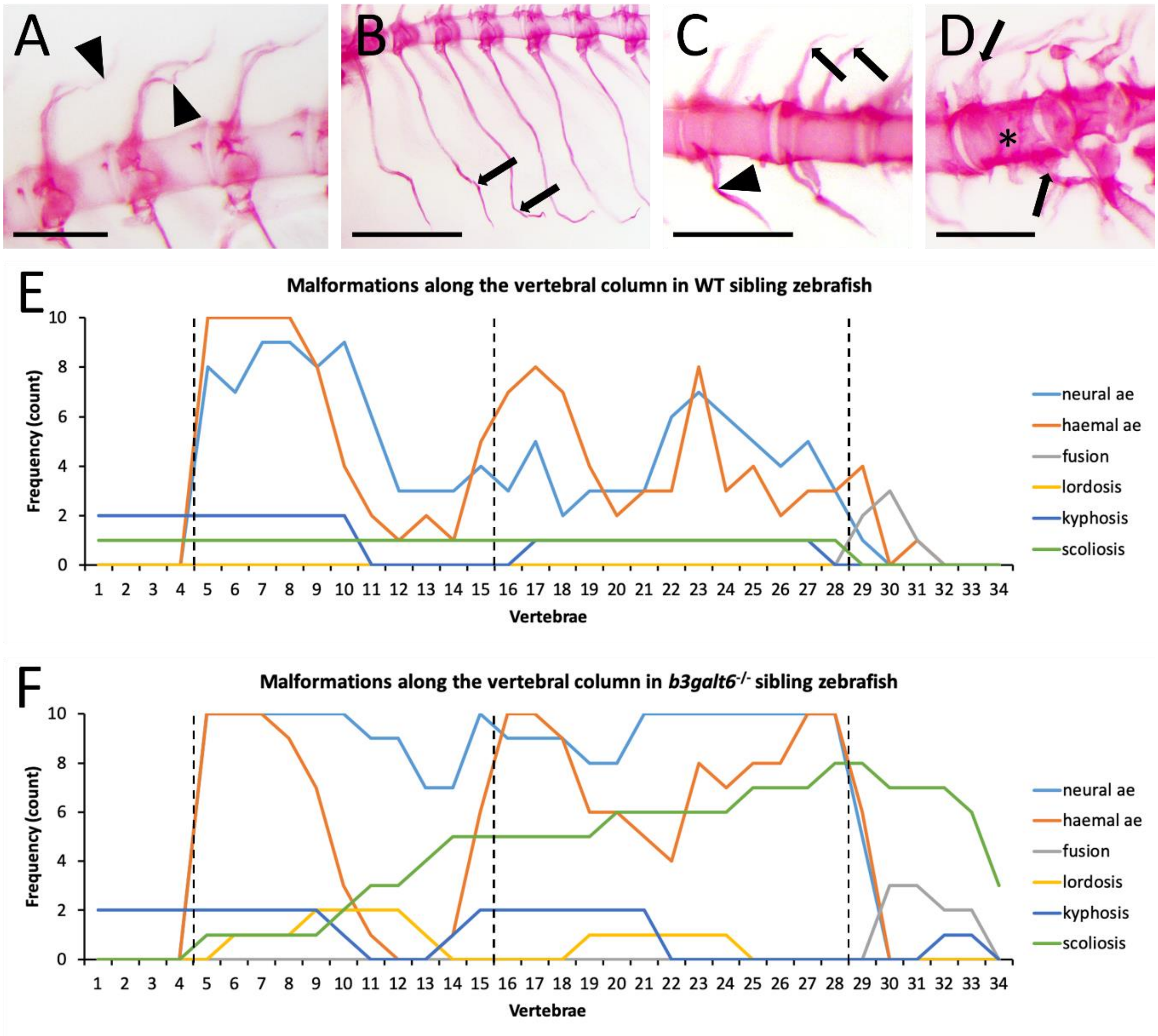


**Fig. S10. Malformations of the vertebral column and their frequency in WT and *b3galt6*$^{-/-}$ siblings at 20 dpf.** All images are oriented anterior to the left, posterior to the right, dorsal to the top and ventral to the bottom. Main malformations in the vertebral column affected the associated elements and included bending of the spine, i.e. lordosis, kyphosis and scoliosis. (**A**) Malformation of the neural associated element with bending of the arch and spine (arrowheads) occurred most frequently. Scale bar: 200 µm (**B**) At the haemal side in the abdominal region bended ribs were observed (block arrows). Scale bar: 500 µm (**C**) At the haemal side in the caudal region mainly bended associated elements were observed (arrowhead). Notice also the unfused neural arches and asymmetric implantation of the neural arches (block arrows). Scale bar: 200 µm (**D**) Vertebral fusion was observed in preural vertebrae (indicated by asterisk) based on the presence of two arches on a fused vertebral centrum. The arrow shows a first neural arch (top of image). While the corresponding haemal arch of the first neural arch was not present, a second haemal arch (indicated by arrows) was present on the same vertebral centrum. Scale bar: 200 µm (**E-F**) The X-axis represents every vertebral position along the vertebral column of the zebrafish. The black dotted lines indicate the borders of the vertebral column regions, the first line indicates the border between the Weberian apparatus and the abdominal region, the second line indicates the border between the abdominal and caudal regions and the third line between the caudal and caudal fin regions. Malformations of the neural and haemal associated elements (ae) includes bending, kinking, lack of fusion, absence and asymmetric implantation. The associated elements of the Weberian apparatus were not assessed for malformations since these elements have specialized functions with no homologues structure in humans and are therefore not relevant for the current research. Vertebral elements of 10 WT and 10 *b3galt6*$^{-/-}$ zebrafish were scored by 2 blinded observers.

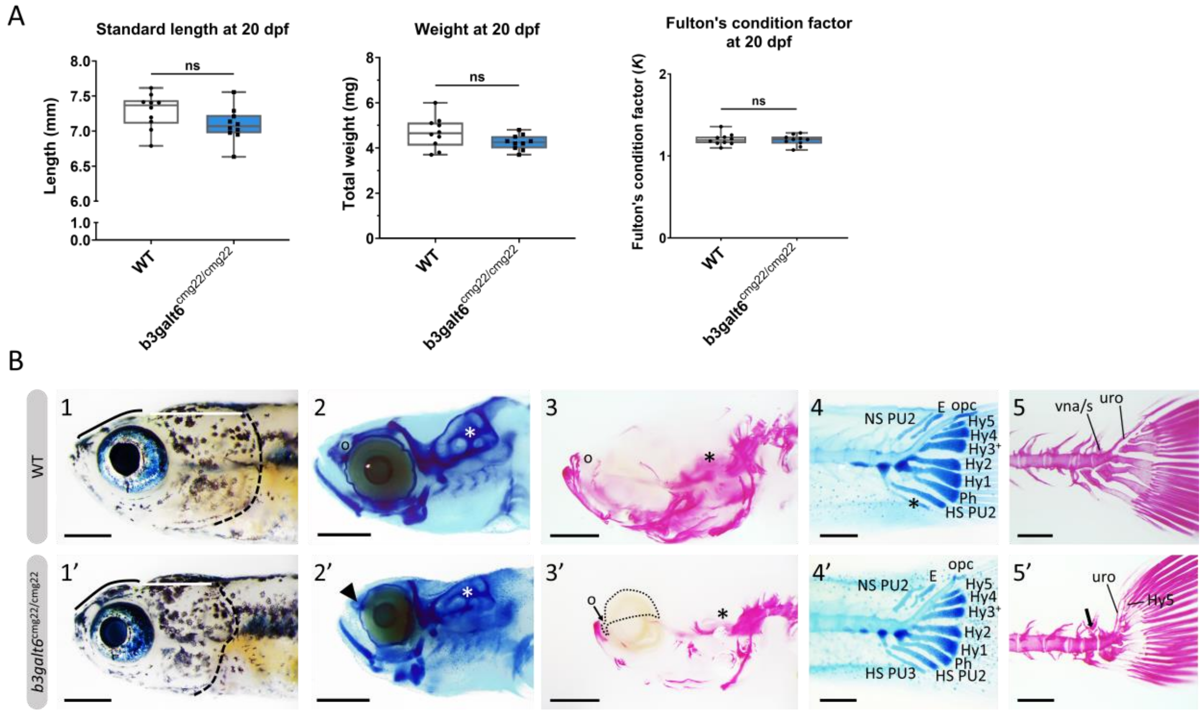


**Fig. S11. External morphological abnormalities and delay of mineralization at 20 dpf in the second *b3galt6*$^{-/-}$ zebrafish harboring cmg22 alleles.** (**A**) Weight, standard length and Fulton's condition factor of juvenile WT and *b3galt6*$^{-/-}$ zebrafish was not significantly different at 20 dpf (n=10). (**B**) All images are oriented anterior to the left, posterior to the right, dorsal to the top and ventral to the bottom. (**1-1'**) External phenotype of the head of juvenile WT and *b3galt6*$^{-/-}$ sibling zebrafish. *b3galt6*$^{-/-}$ zebrafish display a rounder frontal region (black line), shorter parietal/occipital region (white horizontal line) and a malformed posterior edge of the opercular apparatus (dotted black line), compared to WT control. (**2-2'**) Representative images of cartilage AB staining of the head. The olfactory region (indicated by 'o' in the WT and arrowhead in the *b3galt6*$^{-/-}$ image) and the otic region (indicated by white asterisk) barely show delayed development of the cartilage elements in *b3galt6*$^{-/-}$ zebrafish as compared to WT. (**3-3'**) Representative images of AR mineral staining of the head. The olfactory region (indicated by ‘o’) and the otic region (indicated by asterisk) show severely delayed mineralization in *b3galt6*$^{-/-}$ zebrafish juveniles as compared to WT. Also the orbital region showed delayed mineralization (indicated by dotted black line). Note that here the cartilage development and timing of mineralization appear to be uncoupled (equal cartilage development in mutants and WT) compared to the b3galt6$^{-/-}$ cmg20 zebrafish where delayed cartilage developments results in delayed in mineralization. (**4-4’**) Representative images of AB staining of the caudal fin. The modified associated elements show more anomalies, duplications of elements (second HS PU2 indicated by asterisks) and mild alterations in the shape but no delay in development. Abbreviations: E, epural; HS PU2, haemal spine of preural 2; HS PU3, haemal spine of preural 3; Hy1-5, hypural 1-5; NS PU2, neural spine preural 2; opc, opistural cartilage; Ph, parhypural (**5-5’**) Representative images of AR mineral staining of the caudal fin. Caudal vertebrae and their associated elements show almost no delay in mineralization. Malformation or absence of caudal fin arches (block arrow indicates a neural arch), and malformation of uroneural (uro) and vestigial neural arches and spines (vna/s) are observed in *b3galt6*$^{-/-}$ zebrafish; hypurals have a bent appearance. Scale bars: images of the head 500 µm, images of the caudal fin 200 µm.

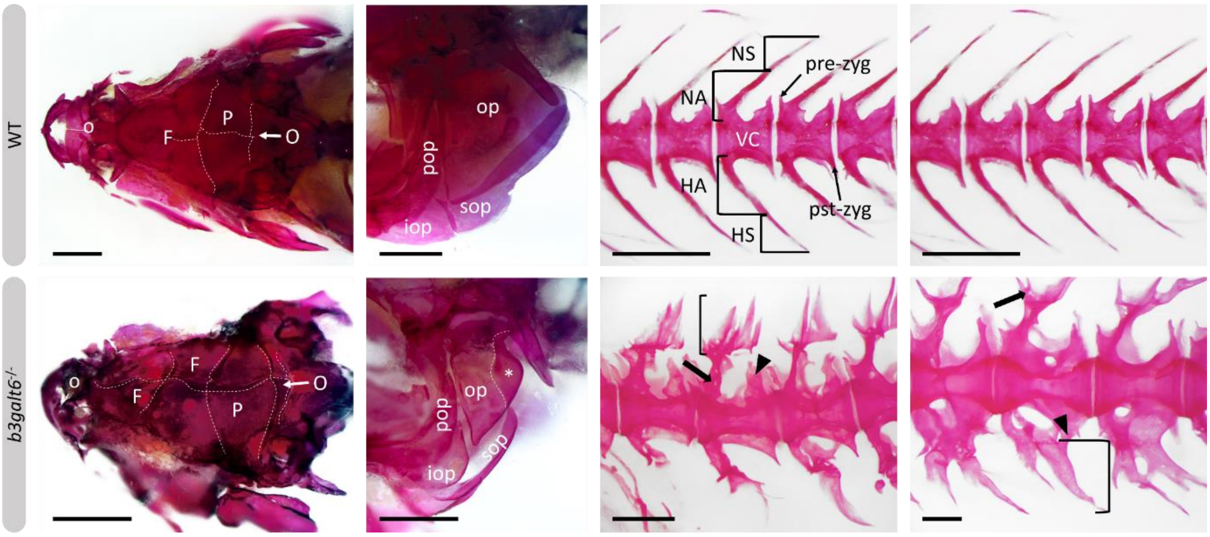


**Fig. S12. Specific malformations of the head and vertebral column in adult *b3galt6*$^{-/-}$ zebrafish.** All images are oriented anterior to the left, posterior to the right, dorsal to the top and ventral to the bottom. At adult stage, developmental or mineralization delay was no longer observed in mutant zebrafish. Dorsal view of the cranial roof bones shows the malformation of the sutures in the cranial roof, with gaps between cranial roof bones rather than overlap. The anterior part of the frontal bone (left F) should no longer show a suture since synostosis occurs when the ossification centra reach the medial line (craniosynostosis as a comparable human term). *b3galt6*$^{-/-}$zebrafish could show a suture in the anterior part of the frontal bone. Notice also the malformed and underdeveloped olfactory region (indicated by 'o') in the *b3galt6*$^{-/-}$ skull. Other abbreviations: O, occipital; P, parietal. Scale bars: 1 mm. While the opercular bone (op) in WT zebrafish had a triangular shape the opercle in severely affected KO zebrafish shows an indentation on the postero-dorsal edge of the bone or the posterior edge of the opercle was backfolded on the opercle itself (indicated by white dotted line and asterisk). Other abbreviations: iop, interopercle, pop, preopercle, sop, subopercle. Scale bars: 1 mm. The two top images of vertebrae show WT caudal vertebrae. The phenotype of abdominal (left image) and caudal (right image) vertebral bodies in KO zebrafish was strikingly different. Abbreviations: HA, haemal arch; HS, haemal spine; NA, neural arch; NS neural spine; pre-zyg, pre-zygapophysis; pst-zyg, post-zygapophysis; VC, vertebral centrum. In the abdominal region the neural arches were often wider (arrow) due to the presence of extra intramembranous bone. Multiple parallel intramembranous spines with flange like outgrowths were present instead of a single neural spine (indicated by bracket). The pre- and post-zygapophyses (arrowhead) on the dorsal and ventral sides extended respectively more dorsal or ventral, were larger and could have spikey intramembranous outgrowths. In the caudal region the neural arch was also wider but only one or two intramembranous neural spines that oriented anteriorly or dorsally were present. Often a third intramembranous spine oriented anteriorly was present where the neural arch fuses to the neural spine (arrow). The wider haemal arches could be fused to a ventrally enlarged and wider post-zygapophysis (arrowhead). On haemal arches fused to the post-zygapophysis at least two haemal spines widened by flange like outgrowths were present (indicated by bracket). Scale bars: 1 mm for WT vertebrae, 500 µm for the abdominal mutant vertebrae and 200 µm for the caudal mutant vertebrae.

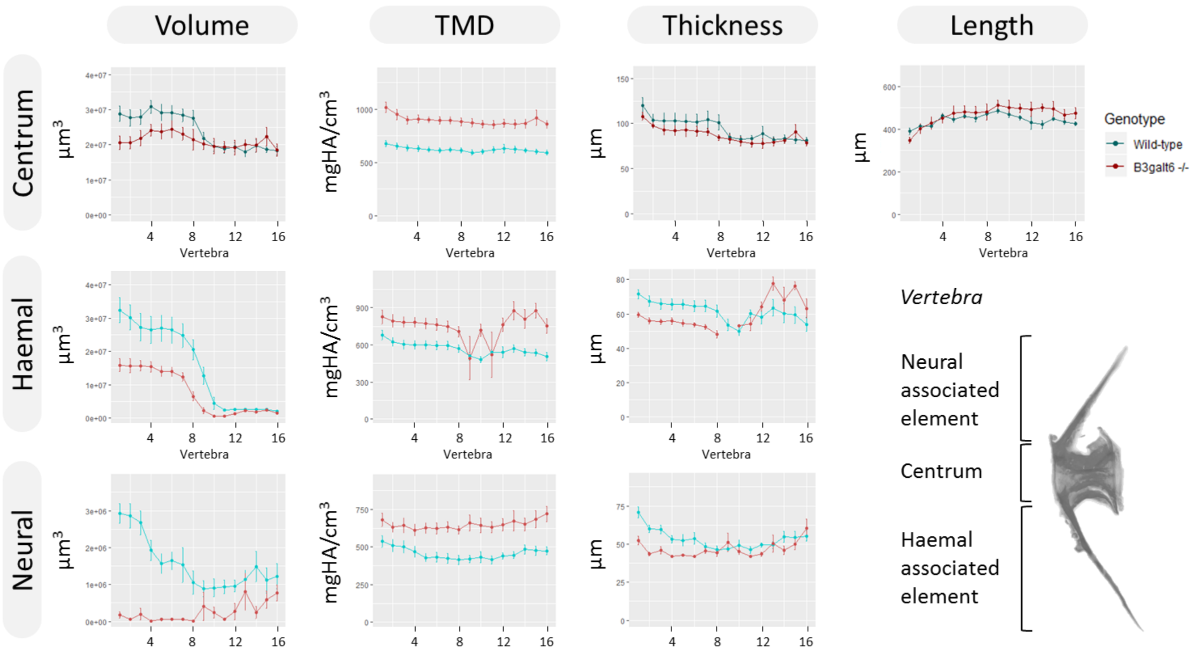

**Fig. S13. µCT analysis of *b3galt6*$^{-/-}$ zebrafish following removal of allometric effects of body size.** Phenotypic features as a function of vertebra in five mutant versus five wild-type zebrafish at the age of four months. Phenotypic data in WT sibling controls were subjected to allometric normalization as outlined by Hur *et al.* (1); data of *b3galt6*$^{-/-}$ zebrafish are unnormalized and thus identical to those in Fig. 6. Plots associated with a significant difference are colored in a lighter coloring scheme. A decrease in volume and thickness of the haemal and neural associated elements is observed after normalization (P=0.01, P=0.000053, P=0.0027 and P=0.0041, respectively) while the centrum volume is not significantly different after normalization (P=0.098). Remarkably, after normalization we clearly note a strong increase of centrum, haemal and neural associated element TMD in our mutants (P=0.00038, P=0.0086 and P=0.0048, respectively). Values for TMD were derived by a two-step process in which tissue mineral concentrate (TMC) and volume were subjected to allometric normalization independently, and normalized values for TMC and volume were used to calculate normalized values for TMD. Data are presented as mean ± SEM. TMD, tissue mineral density; HA, hydroxyapatite.

**Table S9. Hydroxylation status from cross linking sites C-telopeptide and K87 at the α1 chain from type I collagen.** Hyl: hydroxylation, C-telo: C-terminal telopeptide, K87: lysine at position 87. A two-tailed t-test was used to determine the standard error of the mean (SEM).

| Bone sample | % Hyl of α1(I) C-telo | % Hyl of α1(I) K87 | Average % Hyl α1(I) C-telo ± SEM |
|---|---|---|---|
| WT 1 | 70 | 100% Hyl Galactose-Glucose | |
| WT 2 | 79 | 100% Hyl Galactose-Glucose | |
| WT 3 | 60 | 100% Hyl Galactose-Glucose | 70.8 ± 4.5% |
| WT 4 | 83 | 100% Hyl Galactose-Glucose | |
| WT 5 | 62 | 100% Hyl Galactose-Glucose | |
| *b3galt6*$^{-/-}$ 1 | 76 | 100% Hyl Galactose-Glucose | |
| *b3galt6*$^{-/-}$ 2 | 65 | 100% Hyl Galactose-Glucose | |
| *b3galt6*$^{-/-}$ 3 | 70 | 100% Hyl Galactose-Glucose | 68.6 ± 3.3% |
| *b3galt6*$^{-/-}$ 4 | 58 | 100% Hyl Galactose-Glucose | |
| *b3galt6*$^{-/-}$ 5 | 74 | 100% Hyl Galactose-Glucose | |

### Supplementary Material and Methods

Zebrafish Maintenance

Wild-type (AB strain) and *b3galt6* knock-out zebrafish lines were reared and maintained by standard protocols. All zebrafish were housed in ZebTEC semi-closed recirculation housing systems (Techniplast, Italy) and kept at a constant temperature (27–28°C), pH (7.5) and conductivity (500 µS) on a 14/10 light/dark cycle. Zebrafish were fed three times a day with dry food (Gemma Micro, Skretting, Preston, UK) and Artemia (Ocean Nutrition, Essen, Belgium). All animal studies were performed in agreement with EU Directive 2010/63/EU for animals, permit number: ECD 16/18. All zebrafish used in a single experiment were bred at the same density. All efforts were made to minimize pain, distress and discomfort.

CRISPR/Cas9 injections

*b3galt6* CRISPR target sequences (available on request) were identified with CRISPRdirect and CRISPRscan. Cas9-NLS protein (250 pg; LabOmics) and *b3galt6* sgRNA (25 pg) were co-injected into one-cell-stage wild-type (AB) embryos.

RT-qPCR

To compare the transcription levels of *xylt1*, *xylt2*, *b4galt7*, *b3galt6*, *b3gat3*, *b3galt1a*, *b3galt2*, *b3galt4* and *b3galnt2* in six-month-old WT and *b3galt6*$^{-/-}$ adult zebrafish, quantitative reverse-transcription PCR (RT-qPCR) was performed as described before (2) on the caudal fin of WT (n=5) and *b3galt6* KO (n=5) zebrafish. qbase+ 3.0 (Biogazelle, Zwijnaarde, Belgium) was used for data analysis, using reference genes *bactin2*, *elfa*, and ERE elements *hatn10*, *loopern4* and *tdr7* for normalization (3). Data are presented as mean ± standard error of the mean (SEM). Primer sequences are available upon request.

Alcian blue staining

Cartilage was stained with Alcian blue, via a modified protocol of Neuhauss *et al.*, 1996 (4). Wild-type (n=5), and *b3galt6*$^{-/-}$ (n=5) larvae were collected at 20 dpf and fixed overnight in 4% paraformaldehyde (PFA) and washed several times in PBS with 0.1% Tween-20 (PBST). In order to enhance their optical clarity, the specimens were bleached in a solution containing 3% hydrogen peroxide and 2% potassium hydroxide for 2 hr or until the larvae were sufficiently translucent. The larvae were rinsed in PBST and transferred into an Alcian blue solution (3.7% concentrated hydrochloric acid, 70% ethanol, 0.1% Alcian blue) and stained overnight. Specimens were cleared in acidic ethanol (5% concentrated hydrochloric acid, 70% ethanol) for 4 hr and rehydrated in an ethanol series. To further reduce background staining and interference of surrounding tissues, a digest was done with a 1 mg/ml trypsin solution in 60% saturated borax for 15 min. Finally, the specimens were rinsed with demineralized water and gradually transferred to pure glycerol for documentation and storage at 4°C. Analysis was done using a Leica M165 FC Fluorescent Stereo Microscope and a Leica DFC 450C camera (Leica Microsystems, GmbH, Wetzlar, Germany) with LAS V4.3 software.

<u>Alizarin red staining</u>

**Larvae -** Mineralized bone of fixed specimens was stained using a modified protocol (5). WT (n=10) and *b3galt6*$^{-/-}$ (n=10) zebrafish larvae were fixed in 4% PFA in 0.1 M sodium phosphate buffer for 1 hr, washed with demineralized water and subsequently bleached in 1% hydrogen peroxide, 1% potassium hydroxide and 2% Triton X-100 for 30 min. After bleaching, embryos were washed with demineralized water and stained with 0.5% Alizarin red S (C.I. 58005 Sigma) in a 1% potassium hydroxide and 2% Triton X-100 solution for 3 hr. Specimens were cleared for 30 minutes in a 20% glycerol and 0.25% potassium hydroxide solution. Specimens were stored at 4°C after gradual transfer to 100% glycerol. Specimens were imaged using a Leica M165 FC Fluorescent Stereo Microscope and a Leica DFC 450C camera (Leica Microsystems GmbH, Wetzlar, Germany) with LAS V4.3 software.

**Adults -** Five-month-old WT (n=5) and *b3galt6*$^{-/-}$ (n=5) zebrafish were fixed with 4% paraformaldehyde (PFA) for 24 hr at room temperature while gently shaking, washed with demineralized water overnight (O/N) and subsequently bleached in 3% hydrogen peroxide and 1% potassium hydroxide for 4 hr. After bleaching, adults were washed with demineralized water for 30 min and immersed in a 30% saturated borax solution for 15 hr. The specimens were washed three times and stained with 1% Alizarin red S (C.I. 58005 Sigma) in a 1% potassium hydroxide solution for 72 h. Clearing was carried out for 30 minutes in demineralized water and afterwards in a 20% glycerol and 0.1% potassium hydroxide solution for 24 hr. To further reduce background staining, a tissue digest with a 1% trypsin solution in 2% saturated borax was applied O/N while gently shaking. Specimens were stored at 4°C after gradual transfer to 100% glycerol. Specimens were imaged using a Leica M165 FC Fluorescent Stereo Microscope and a Leica DFC 450C camera (Leica Microsystems GmbH, Wetzlar, Germany).

Terminology of cartilaginous and bony skeletal structures was adapted from Cubbage and Mabee (6) for the cranium, Dewit *et al.* (7) for the pectoral fin, Arratia *et al.* (8) for the vertebrae, De Clercq *et al.* (9) for vertebral column regions, and Bensimon-Brito *et al.* (10) for the caudal fin. Types of bone were defined based on the mode of ossification following Hall (11). Vertebral deformity types were classified based on Witten *et al.* (12). All samples were oriented with anterior to the left and dorsal to the top. Development of cartilaginous structures was assessed based on a dorsal view and a lateral view of the left side. Mineralization status was determined on bones on the lefts side of the specimen unless for median bones where either a dorsal or ventral view was used. If the observations were unclear on the left side, the representative bone on the right side of the specimens was used.

<u>µCT scanning analysis</u>

For µCT-based phenotyping and quantification, four-month-old adult zebrafish were euthanized using 0.4% tricaine, fixed in 4% PFA for 48 hours and transferred to a 70% ethanol solution for scanning. Whole-body µCT scans of WT (n=5) and *b3galt6*$^{-/-}$ (n=5) siblings were acquired on a SkyScan 1275 (Bruker, Kontich, Belgium) using the following scan parameters: 0.25 mm aluminum filter, 50 kV, 160 µA, 65 ms integration time, 0.5° rotation step, 721 projections/360°, 5 m 57 s scan duration and 21 µm voxel size. Two zebrafish were scanned simultaneously in each acquisition and DICOM files of individual fish were generated using NRecon V1.7.3.2 (Bruker) software. For each zebrafish, standard length (the length from the tip of the snout to the posterior end of the last vertebra) was measured on maximal projections of DICOM files via ImageJ 1.52i. DICOM files of individual zebrafish were segmented in MATLAB using custom FishCuT software and data were analyzed and corrected for standard length in the R statistical environment, as previously described (1). The global test was used to assess whether the pattern across vertebrae was significantly different between a certain mutant genotype and its control group of WT siblings.

<u>Histology and transmission electron microscopy</u>

Zebrafish bone, consisting of the last transitional vertebra and two consecutive caudal vertebrae (13, 14), skin, at the lateral line below the dorsal fin, and muscle samples, located dorsolateral underneath the dorsal fin, dissected from five-month-old WT (n=5) and *b3galt6*$^{-/-}$ (n=5) zebrafish, were fixed and embedded in epon following the procedures outlined by Huysseune and Sire (15). All semi-thin (500 nm) sections were stained with toluidine blue and examined using a Zeiss Axio Imager Microscope and photographed using an Axiocam MRC camera. Ultra-thin (80 nm) parasaggital sections were cut using a Leica Ultracut UC7 ultramicrotome with a diamond knife (Diatome) and collected on formvar-coated copper single slot grids (Agar Scientific). Sections were studied and photographed with a JEOL JEM1010 TEM (Jeol Ltd., Tokyo) equipped with a CCD side mounted Veleta camera. Pictures were digitized using a Ditabis system (Ditabis Ltd., Pforzheim, Germany). Resulting images were processed with Fiji software (16).

Swim tunnel experiments

Ten wild-type (five months) and ten *b3galt6*$^{-/-}$ zebrafish (eight months), were selected based on their similar (not significantly different) length and weight.

The critical swimming speed ($U_{crit}$) was measured for both WT and *b3galt6*$^{-/-}$ zebrafish using a swim tunnel. Critical swimming speeds were calculated using the equation: $U_{\mathrm{crit}} = U_{\mathrm{i}} + [U_{\mathrm{ii}}(T_{\mathrm{i}}/T_{\mathrm{ii}})]$, where $U_{\mathrm{i}}$ is the highest velocity maintained for the whole 5 min (cm/s), $U_{\mathrm{ii}}$ is the velocity increment (5 cm/s), $T_{\mathrm{i}}$ is the time elapsed at fatigue velocity and $T_{\mathrm{ii}}$ is the time between velocity changes (5 min) (17). In short, each zebrafish was allowed a one minute warm up at a velocity of 7.5 cm/s, thereafter the zebrafish were swimming for five minutes at the following speeds: 18, 25, 30, 35, 40, 45, 50 and 55 cm/s. When the zebrafish were not able to swim anymore, the test was stopped and the time they lasted determined the $U_{crit.}$.

Furthermore, the enducrance of each zebrafish was tested by means of an effort test in the swim tunnel. Each zebrafish swum 1 minute at a speed of 15 cm/s, the seconde minute at 24 cm/s and the third minute at 28 cm/s. After the third minute, the speed increased until 37 cm/s for maximum 7 minutes. When the zebrafish was not able to swim anymore, the test was stopped. In this way, the stamina of each zebrafish was determined.

Disaccharide composition analysis of CS, DS, and HS chains isolated from dissected zebrafish tissues

The level of total disaccharides from CS, DS, and HS in nine-month-old zebrafish were determined as described previously (18, 19). Briefly, bone (vertebral column, excluding Weberian apparatus and urostyle), muscle, and skin of wild-type (n=3) and *b3galt6*$^{-/-}$ (n=3) zebrafish were collected, sonicated, and treated exhaustively with actinase E (Kaken Pharm., Kyoto, Japan) to remove proteins. GAG fractions were obtained by precipitation with 80% ethanol containing sodium acetate and desalted with an Amicon Ultra-4. The samples were treated individually with a mixture of chondroitinases ABC and AC-II (Seikagaku Corp., Tokyo, Japan), a mixture of chondroitinases AC-I and AC-II, chondroitinase

B (IBEX Technologies, Montreal, Canada), or a mixture of heparinase-I and heparinase-III (IBEX Technologies) for analyzing the disaccharide composition of CS/DS, CS, DS or HS moieties, respectively. The digests were labeled with a fluorophore 2-aminobenzamide (2AB) and aliquots of the 2AB-derivatives of CS/DS/HS disaccharides were analyzed by anion-exchange HPLC on a PA-G column (YMC Co., Kyoto, Japan) as described previously (19, 20). The unsaturated CS/DS or HS disaccharides observed in the digests were identified by comparison with the elution positions of authentic 2AB-labeled disaccharide standards.

Glycopeptide preparation and analysis

**LC-MS/MS sample preparation -** CS- and HS-glycopeptides were purified from six-month-old WT (n=3) and *b3galt6*$^{-/-}$ (n=3) zebrafish using a combination of extraction, trypsin digestion and anion exchange chromatography, modified from previously described protocols (21, 22). In brief, one zebrafish was prepared at a time (WT or KO) by homogenizing the zebrafish in 1 mL, 1% CHAPS buffer using a Dounce homogenizer. The samples were boiled for 10 min at 96 °C and approximately 0.5 mg of each homogenate was then reduced and alkylated in 50 mM $NH_4HCO_3$ in the presence of Protease Max surfactant trypsin enhancer (0.02% final concentration) (Promega). Additional Protease Max surfactant was thereafter added (0.03% final concentration) and the samples were trypsinized over night (37°C) with 40 µg trypsin (Promega). The trypsin-digested sample was enriched for GAG-peptides using SAX-chromatography (Vivapure, Q Maxi H). The digests were diluted in 50 mL of 50 mM sodium acetate, 0.2 M NaCl, pH 4.0 and the material was then applied onto the columns. The column was spun for 1500 x g for 2 min, and the procedure was repeated until all material had been applied onto the column. The columns were then washed with 20 mL 50 mM Tris-HCl, 0.2 M NaCl, pH 8.0. The GAG-peptides were eluted in a single step using 50 mM Tris-HCl, pH 8.0, 2.0 M NaCl. The fractions were desalted using PD-10 columns, lyophilized and treated with Chondroitinase ABC (C3669, Sigma Aldrich) and heparinase II and III (EC 4.2.2.8) (overexpressed in E.coli, gift from Prof. Jian Liu), respectively. The lyophilized material was dissolved in 100 µL of 50 mM $NH_4Ac$, 4 mM $CaCl_2$ pH 7.3 and then 10 mU of Chondroitinase ABC, heparinase II and III were added to the sample and incubated for 37°C for 12 h. Prior to the LC-MS/MS -analysis the samples were desalted with C18 spin columns (Thermo Fisher Scientific) and dried.

**LC-MS/MS analysis -** The samples were analyzed on an Orbitrap Fusion mass spectrometer coupled to an Easy-nLC 1200 liquid chromatography system (Thermo Fisher Scientific., Waltham, MA). Glycopeptides (3 µL injection volume) were trapped on an Acclaim Pepmap 100 C18 trap column (100 µm x 2 cm, particle size 5 µm, Thermo Fischer Scientific) and separated on an analytical column (75 µm x 30 cm) packed in-house with Reprosil-Pur C18 material (particle size 3 µm, Dr. Maisch, Germany). The following gradient was run at 300 nL/min; 7–35% B-solvent over 45 min, 35–100% B over 5 min, with a final hold at 100% B for 10 min; solvent A was 0.2% formic acid (FA) in water, solvent B was 80% acetonitrile, 0.2% FA.

Nanospray Flex ion source was operated in positive ionization mode at 1.8 kV. MS scans were performed at 120,000 resolution (at m/z 200), with an Automatic Gain Control (AGC)-target value of 1E6 and scan range of m/z 600–2000. The most abundant precursor ions with charges from +2 to +7 were selected for fragmentation with the maximum cycle time of 3 s and the dynamic exclusion with a duration of 10 s. Three separate higher-energy collision-induced dissociation (HCD) MS/MS spectra were recorded for each precursor, with scan 1 at the HCD collision energy of 30% (NCE) with a scan range from 100 to 2000, scan 2 at the HCD energy of 40% with a scan range from 300 to 2000 and scan 3 at the HCD energy of 40% with the scan range from 100 to 2000. All MS/MS scans were acquired with the precursor isolation window of 5 Th, resolution 30,000 (at *m/z* 200), AGC target of 1e5 and maximum injection time of 118 ms.

**LC-MS/MS data analysis -** The glycopeptide files were analyzed using both manual interpretation and automated Mascot searches for GAG-glycopeptides. The files were manually interpreted using the Xcalibur software (Thermo Fisher Scientific). Database searches were performed against *Danio rerio* in the UniProtKB/Swiss-Prot database and NCBI using Mascot Distiller (version 2.6.1.0, Matrix Science, London, U.K) and an in-house Mascot server (version 2.5.1). The Mascot searches employed the criteria *Trypsin* as enzyme specificity, allowing for up to two missed cleavages. The peptide tolerance was 5 ppm and the fragment tolerance 20 ppm. The searches included variable linkage region structures at serine residues, including both hexa- and pentasaccharide modifications, as predicted after chondroitinase ABC degradation. The hexasaccharide modifications included the hexasaccharide structure [HexA(-$H_2O$)-HexNAc-HexA-Hex-Hex-Xyl-O-] with 0 ($C_{37}H_{55}NO_{30}$, 993.2809 Da), 1 ($C_{37}H_{55}NO_{33}S$, 1073.2377 Da), or 2 ($C_{37}H_{55}NO_{36}S_2$, 1153.1945 Da) sulfate groups attached. The pentasaccharide modifications included the pentasaccharide structure [HexA(-$H_2O$)-HexNAc-HexA-Hex-Xyl-O-] with 0 ($C_{31}H_{45}NO_{25}$, 831.2281 u), 1 ($C_{31}H_{45}NO_{28}S$, 911.1849 u), or 2 ($C_{31}H_{45}NO_{31}S_2$, 991.1417 u) sulfate groups, including neutral loss of the corresponding glycan modifications.

It should be noted that some glycan structures may have phosphate- (79.9568 Da) instead of sulfate modifications (79.9663 Da). Based on the small mass difference, the distinction of sulfate- and phosphate modification cannot be made only on the MS1 precursor weight, but has to be manually evaluated also at the MS2- level.

GlcAT-I assay

GlcA-transferase assay was carried out by the methodology as previously described (23-25), with slight modifications. Briefly, UDP-Glo Glycosyltransferase detection kit (Promega), was utilized for the assay. The recombinant B3GAT3/GlcAT-I was transiently expressed in HEK293T cells using p3xFLAG-CMV8-B3GAT3 and Lipofectamine 3000. Furthermore, Galβ1-3Galβ1-*O*-methyl (Sigma, St. Louis, MO, USA) and Galβ1-4Xylβ-*p*-nitrophenyl (Medical Chemistry Pharmaceutical, Sapporo, Japan), were used as sugar acceptors for the B3GAT3/GlcAT-I assay.
